# Low anabolic independence emerges when cultivating more than three bacterial species together

**DOI:** 10.1101/2025.04.28.650956

**Authors:** Armando Pacheco-Valenciana, Anna Tausch, Iva Veseli, Jennah E. Dharamshi, Fabian Bergland, Luis Fernando Delgado-Zambrano, Alejandro Rodríguez-Gijón, Anders F. Andersson, Sarahi L. Garcia

## Abstract

Microorganisms thrive in complex communities shaped by intricate interactions, yet the extent and ecological implications of biosynthetic dependencies in natural communities remain underexplored. Here, we used a dilution approach to cultivate 204 microbial model communities from the Baltic Sea and recovered 527 metagenome-assembled genomes (MAGs) that dereplicated into 72 species-clusters (>95% average nucleotide identity - ANI). Of these species, at least 70% represent previously uncultivated lineages. Combined with 1073 MAGs from Baltic Sea metagenomes, we generated a genomic catalog of 701 species-clusters. Our results show that cultures with more than 3 species included microorganisms with smaller genome sizes, lower biosynthetic potential for amino acids and B vitamins, and higher prevalence and abundance in the environment. Moreover, biosynthetic genes were distributed across genomes within the same model community, indicating metabolic interdependencies. Our results demonstrate that cultivating bacteria in dilution model communities facilitates access to previously uncultivated but abundant species that likely depend on metabolic partners for survival. Together, our findings highlight the value of community-based cultivation for unraveling ecological strategies. Finally, we confirm that metabolic interdependencies and genome streamlining are widespread features of successful environmental microorganisms.

## INTRODUCTION

Microbial communities in diverse environments operate as complex systems driven by multi-species interactions^1^. Understanding such complex interactions is essential because microorganisms play key roles in the biogeochemical cycles on Earth^2^. To unravel microbial interactions, we need to investigate microbial communities at various levels of biological organization^3,4^, ranging from one-to-one species interactions to simplified multi-species systems (e.g., model communities^5^, synthetic communities^6^, or microcosms^7^), and ultimately to naturally occurring communities.

At the community level, metagenomics has become a powerful tool for uncovering the genetic potential of microbial communities via shotgun sequencing^8–10^. Analyzing metagenomic data reveals not only the vast diversity of microbial species^11^ but also their metabolic potential and co-occurrence networks, which are important for ecosystem functioning^12^. To bridge the gap between broad metagenomic insights and detailed ecological understanding, a few studies have explored genome-specific traits and potential interactions by inferring auxotrophies. While auxotrophs have historically been experimentally identified via cultures that require the addition of specific nutrients to grow^13^, recent work based on genomes and metagenomes has determined auxotrophies based on pathway completeness and found smaller genomes to be more auxotrophic^6,14–19^. In all these studies, auxotrophy has been treated as a binary trait, however, microbial biosynthetic capabilities in nature likely span a spectrum. For example, many microorganisms can complete the biosynthesis of an essential metabolite starting from a precursor or intermediate without needing the essential metabolite itself^20,21^. Nevertheless, modeling work has shown that microbial communities enriched in the so-called auxotrophs can exhibit greater robustness under ecological disturbances, suggesting that these metabolic interdependencies may contribute to overall community stability^22^. While metagenomics offers a broad understanding of microbial communities, interactions cannot easily be inferred from co-occurrences within natural complex ecosystems.

Experimental systems are needed to observe microbial interaction dynamics under controlled conditions. Studies have increasingly turned to simplified systems^23–25^, such as co-cultures^26–28^ and mixed cultures^18,29,30^, in order to identify specific types of interactions. To further contextualize these findings, our literature review in **Table S1** provides a comprehensive overview of publications where microbial interaction patterns were observed in different experimental settings. Across these studies, cross-feeding mechanisms and mutualistic interactions are mostly studied in cultures with two different populations or species^27,31,32^. To a lesser extent, more complex metabolic interactions have also been studied by mixing different isolated species or co-cultivating them in model ecosystems, with the goal of increasing complexity to more closely resemble natural environments^29,33,34^. However, many of these methods focus on cultured isolates, and the vast majority of microorganisms remain uncultivated^35^. An alternative, yet underutilized method of establishing model ecosystems composed of previously uncultivated microorganisms is through dilution cultivation. Dilution to mixed-cultures of naturally co-occurring microorganisms has the potential to cultivate previously uncultivated microorganisms as well as to allow observation of natural microbial interactions^36^. Such cultures, also known as microbial model communities, represent a small subset of the many interactions likely occurring in the natural system^33^. By studying a larger number of microbial model communities, we can gain a more comprehensive understanding of microbial interactions occurring in a natural environment.

In our study, we have focused on studying potential interdependencies at two levels of biological organization by using high-throughput dilution cultivation of model communities together with genome-resolved metagenomics to unravel ecological strategies of microorganisms in the Baltic Sea. Moreover, we examine the biosynthesis of essential metabolites or anabolic independence as a continuous spectrum rather than through conventional binary classifications. For this, we use pathway completeness metrics rather than assigning genomes as strictly prototrophic or auxotrophic. Finally, we identified correlations between genome size, potential biosynthesis of essential metabolites, relative abundance, and prevalence using genomes obtained from both microbial model communities and metagenomic data from Baltic Sea pelagic samples. Our findings demonstrate that microbial model communities are an effective cultivation technique to cultivate previously uncultivated taxa and for identifying putative microbial interactions, including metabolic interdependencies between biosynthetically dependent members.

## RESULTS

### A Baltic Sea MAG catalog

To generate the microbial model communities, we used the dilution-to-extinction cultivation technique in two formats. The first type, low inoculum size, involved inoculating between approximately 2 and 100 cells per well in 1 ml 96-well plates used for each of the inoculum sizes. The second type, high inoculum size, ranged from approximately 200 to 1×10^6^ cells inoculated per microbial model community in 100 ml volumes. After a 4-week incubation period, we sent an aliquot of all 801 cultures for lysis and DNA amplification using multiple displacement amplification (MDA). Based on amplification success, 315 cultures passed the negative control threshold and were sent for sequencing. In total, only 204 microbial model communities together yielded 527 MAGs. Moreover, from the original sample used to establish the microbial model communities, we generated two metagenomes (each from a distinct DNA extraction method) that yielded 305 MAGs. To create a comprehensive genomic catalog, we also added 771 MAGs from 110 publicly available Baltic Sea metagenomes (**Figure 1A**, **Table S2**)^37–39^.

**Figure 1.**
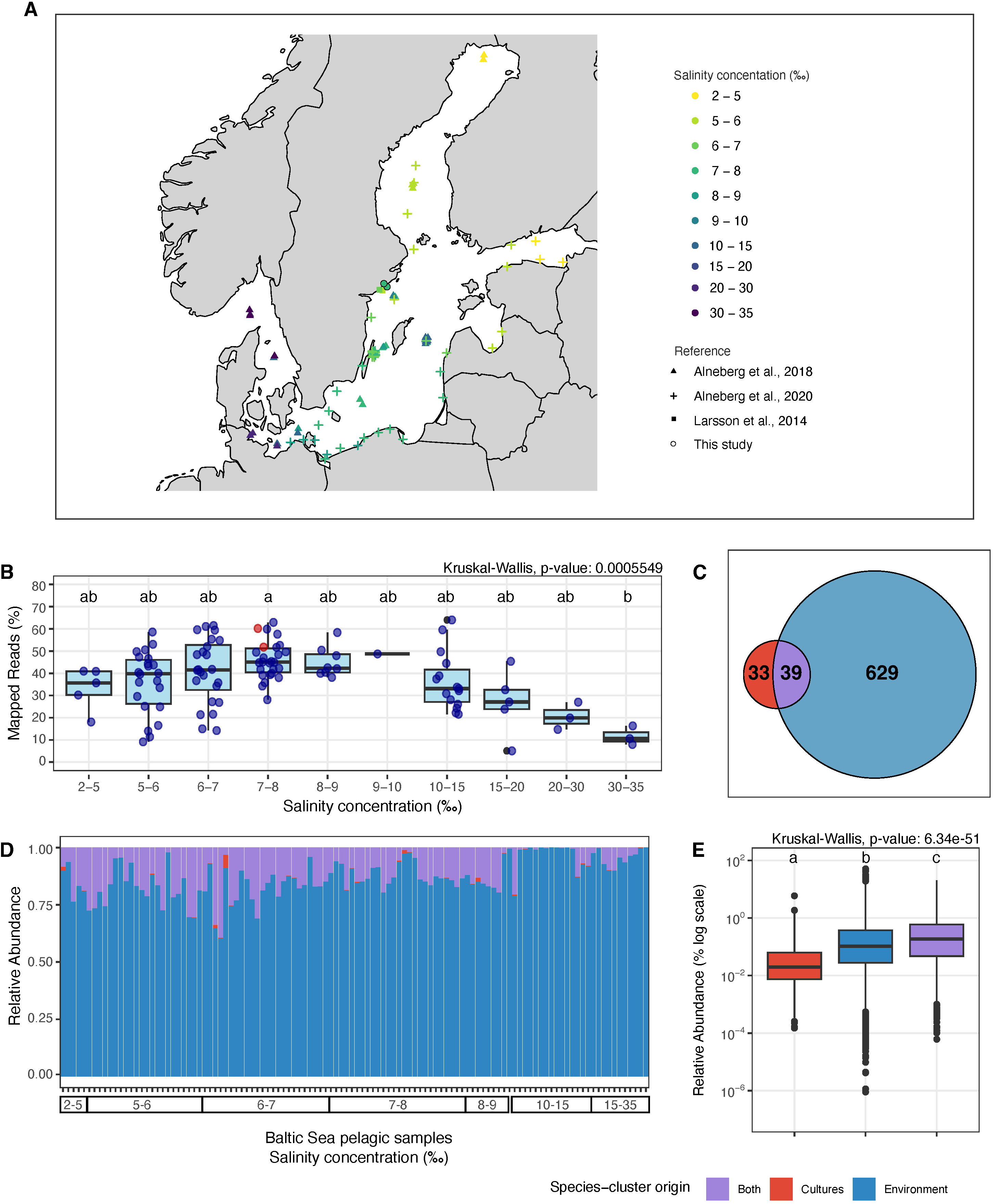
The BalticMAG catalogue. (A) Map of the Baltic Sea showing the geographic location of all the metagenomes analyzed in this study, including our sampling site (n =112). Sampling sites are color-coded according to their salinity gradient (measured in ‰, equivalent to PSU or parts per thousand). The shapes correspond to the different reference sources. (B) Boxplot showing the percentage of mapped reads from all metagenomes to our species-cluster collection, categorized by salinity concentration (Kruskal-Wallis test). (C) Venn diagram showing the overlap of species-clusters presence among cultures (red), the environment (blue), and found in both (purple). (D) Relative abundance of species-clusters across the collection of metagenomes, ordered by salinity gradient from left to right (corresponding to the order in panel B). (E) Boxplot comparing the relative abundance of species-clusters categorized by their presence in cultures, the environment, or both (Kruskal-Wallis test). Kruskal-Wallis tests were followed by Dunn’s post hoc test for pairwise comparisons. Groups sharing at least one letter (e.g., a and ab) are not significantly different from each other, while groups with different letters (e.g., a vs. b) are significantly different (p < 0.05).

Examining this comprehensive MAG catalog allowed us to assess microbial genomic characteristics of all microorganisms found both in the environmental samples and our 204 microbial model communities (**Table S3**). Altogether, the 1603 MAGs were dereplicated into 701 species-clusters (ANI >95%), which form the Baltic Sea genomic catalog (BalticMAG catalog)^40^ used in this study (**Table S4**). The average completeness of the 701 species-cluster representative MAGs is 88%, and they were all used to analyze taxonomy, abundance, prevalence, and estimated genome size (**Figure S1**). The varying completeness of the MAGs has a very minor effect on estimated genome size and relative abundance, just as observed in other studies^17,41^. However, only 450 species-cluster representative MAGs are of high-quality (completeness >90% and contamination <5%) and were used to investigate anabolic potential (**Figure S1D)**.

Examining the source of genomes in the BalticMAG catalog, we found that 33 of the species-clusters included MAGs exclusively from the microbial model communities, 629 included MAGs exclusively from the environmental metagenomes, and 39 (54% of all cultured species) included MAGs from both sources (**Figure 1C, Table S4, Figure S2**).

To investigate the relative abundance of the BalticMAG catalog, we mapped all the environmental metagenomics reads against the genome catalog. We observed that salinity significantly co-varied with the proportion of metagenomic reads that mapped to all species representative genomes (**Figure 1B**). While on average, 38.63% of the metagenomic reads per sample mapped to the BalticMAG catalog, the highest mapping percentage (63.99% and 62.89%) was observed at salinity concentrations 11.28‰ and 7.65‰, respectively. Notably, the two metagenomes from this study (sample with salinity concentration 7.12‰) displayed some of the highest mapping rates at 51.77% and 60.16%, reflecting that the BalticMAG catalog is most complete for salinities between 6 and 11‰.

Despite salinity differences (**Figure 1D**), the average relative abundance of the 39 species-clusters that included MAGs from both microbial model communities and environmental metagenomes was significantly higher in the whole dataset (**Figure 1E**), but also when comparing only the metagenomes from the location and salinity from which we sampled **(Figure S3**). This group of species-clusters (from both sources) shows that our cultivation method can capture some of the most abundant taxa from the environment. Altogether, the diverse taxa cultivated in model communities accounted for ∼20% of the total relative abundance in the original environmental sample (**Figure S4**). Moreover, the 33 species-clusters with MAGs sourced exclusively from microbial model communities were detected across environmental metagenomes, albeit at significantly lower abundances. This indicates that our cultivation approach also enables the recovery of species that are missed by assembly and binning in metagenomic surveys.

### Higher inoculum size increases community richness and uncovers a genome size plateau

In the low inoculum size microbial model communities, the more cells we inoculated, the higher the number of cultures that yielded MAGs (**Figure 2A**). In total, 94 low inoculum size model communities yielded only one MAG each, while 110 model communities resulted in two or more MAGs. In model communities with more than one MAG, each MAG belonged to a different species-cluster in our analysis. Therefore, for clarity, we refer to different MAGs within a model community as different species. Starting from an inoculum size of approximately 30 cells, microbial model communities with more than two species appear more often (**Figure 2B** and **C**). Nearly 82% (n = 433) of the microbial model community MAGs were obtained from multi-species cultures, and the highest number of co-occurring species were found in the high inoculum size microbial model communities inoculated with 5000 cells (**Figure 2C**). Despite using a complex inoculum from the Baltic Sea, observing a maximum of 13 co-occurring species suggests that our cultivation conditions may impose a threshold on the complexity of model communities. Alternatively, since sequencing followed MDA, there is also the possibility that some model communities included more species that were not amplified or assembled and binned. Nevertheless, the increased growth success with increasing inoculum size likely reflects a greater probability of including cells that can grow in isolation or in the presence of a specific required community partner.

**Figure 2.**
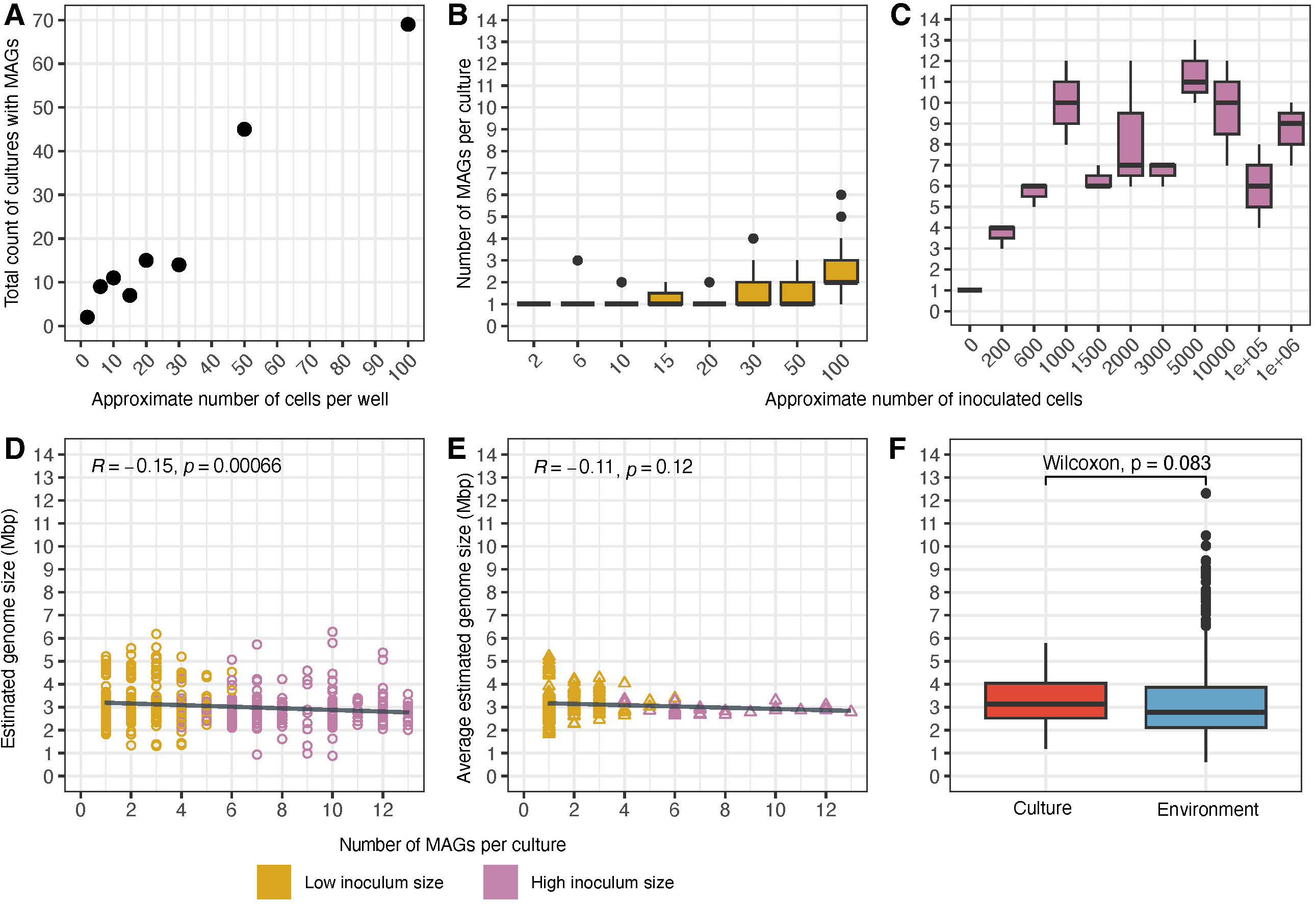
Impact of inoculum size on microbial richness and genome size. (A) Relationship between the number of inoculated cells (low inoculum size = 0 to 100 cells/well) and the total number of cultures from which MAGs were assembled and binned. (B) Number of MAGs per culture for the low inoculum size (yellow) and (C) the high inoculum size (pink) microbial model communities. (D) Estimated genome size (circles) of MAGs from cultures (n = 527) in relation to the number of species growing in the same culture, and (E) average estimated genome size (triangles) of MAGs growing in the same culture. (F) Boxplot comparing the average estimated genome size of species-clusters from cultures and those from the environment (Wilcoxon rank-sum test).

We found that as the number of retrieved MAGs per microbial model community increased, more microorganisms with smaller genomes emerged, as indicated by a slight but significant correlation between estimated genome size and the number of MAGs per microbial model community (**Figure 2D**). In general, the estimated genome sizes ranged from 0.88 to 6.27 Mbp for MAGS from cultures and, from 0.61 to 12.32 Mbp for MAGs from the environment (**Figure 2D** and **F**). While there was no significant difference between the average estimated genome sizes of species-clusters from microbial model communities and Baltic Sea metagenomes (around 3.00 Mbp), we found that species-clusters exclusively composed of genomes from microbial model communities had significantly larger genome sizes than species-clusters including genomes from both sources, and no statistically different genome completeness or contamination (**Figure S5**). Further analysis reinforced the observation that genomes from high inoculum size microbial model communities and from model communities with more than 3 species had on average, smaller genome sizes (2.89 and 2.92 Mbp, respectively - **Figure S6** and **S7**). Genomes from high inoculum size microbial model communities were also on average more prevalent and had higher average relative abundance in the investigated Baltic Sea metagenomes. Finally, the average genome size per microbial model community stabilized at around 3 Mbp in cultures containing more than three species (**Figure 2E**).

### Model communities with more than three species reveal distinct microbial diversity

The 72 species from microbial model communities spanned five of the most abundant phyla (*Pseudomonadota*, *Bacteroidota*, *Campylobacterota*, *Cyanobacteriota*, and *Verrucomicrobiota*) present in the Baltic Sea environmental metagenomic sample^42,43^. These species varied substantially in their observed growth strategies. Four of them were only found in single-species cultures, forty-eight exclusively in multi-species cultures, and twenty appeared growing both alone and in groups (**Figure 3**). Of the species growing consistently in groups, nine grew across different levels of community complexity (e.g., 2, 3, or more than three species per culture), and thirty-nine species were restricted to a single type of community complexity. Specifically, thirty-one species were recovered from microbial model communities with more than three species. Notably, 70% of these cultured species lacked a species-level assignment in the GTDB taxonomy, suggesting they represent previously uncultivated lineages with no characterized MAGs either.

**Figure 3.**
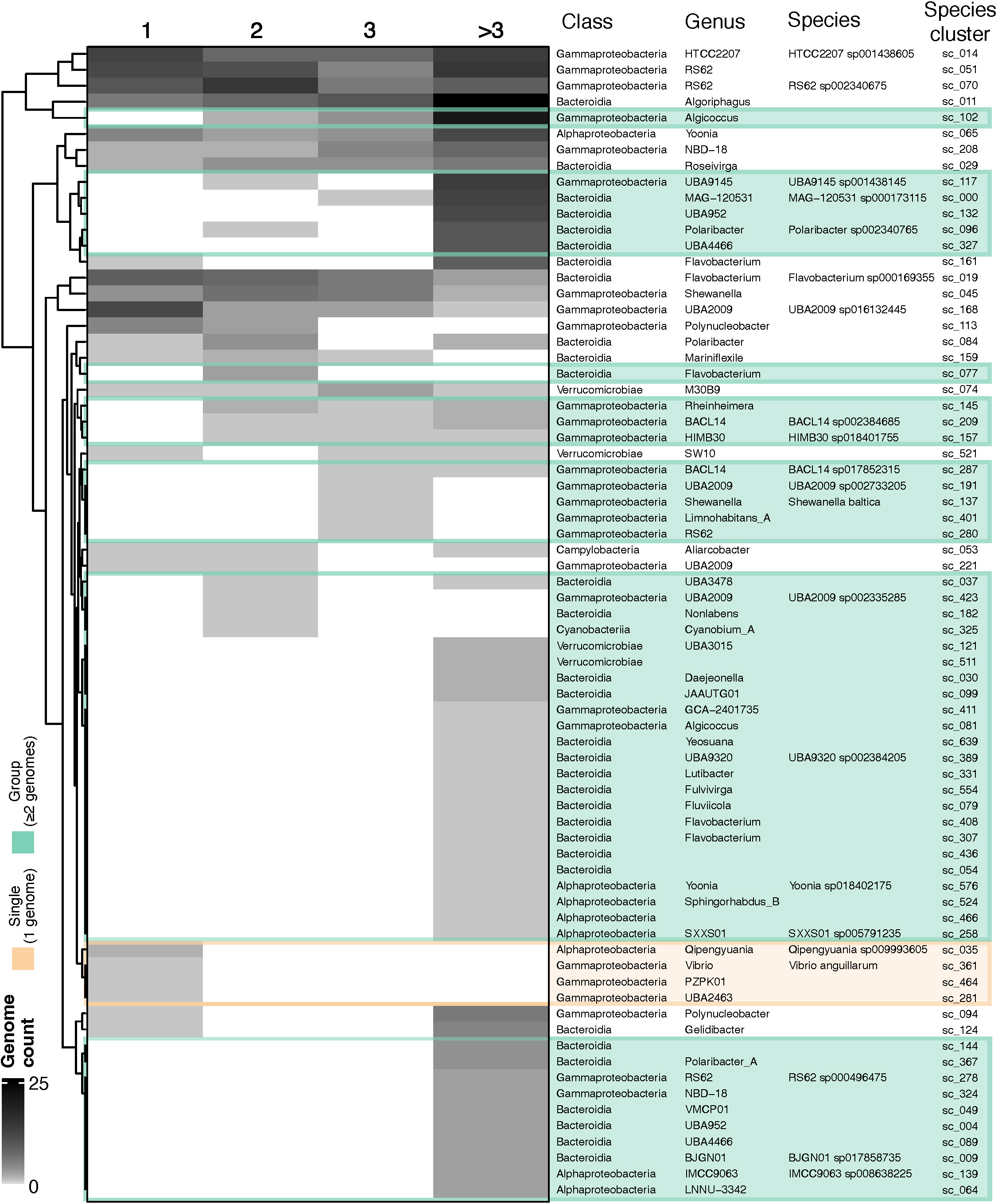
Microbial model communities of increased complexity host distinct sets of cultured species. Heatmap showing the distribution of the 72 species-clusters across cultures grouped by community complexity: 1, 2, 3, or more than 3 species per culture. Taxonomic affiliations (class, genus, and species) based on GTDB-tk are displayed alongside their unique species-cluster IDs. The white-to-black gradient indicates the number of genomes recovered per species-cluster in each culture category. Species that exclusively grow alone are highlighted in a light orange box, while those only found to grow in groups (≥2 genomes) are highlighted in light green.

To evaluate whether culturing bacteria in groups increases the cultivability and recovery of microbial diversity, we computed rarefaction curves for species accumulation across increasing numbers of cultures (**Figure 4**). After 50 cultures of the single-species type, we had in average recovered seventeen species, and every 12 new cultures would yield only three more species in average. As we cultivated more species together, the initial steep slope also increased. In fact, in 25 cultures with more than three species, forty-nine species were recovered. This demonstrates that culturing in groups is a powerful strategy for the recovering of microbial taxa that would normally not grow in axenic cultures.

**Figure 4.**
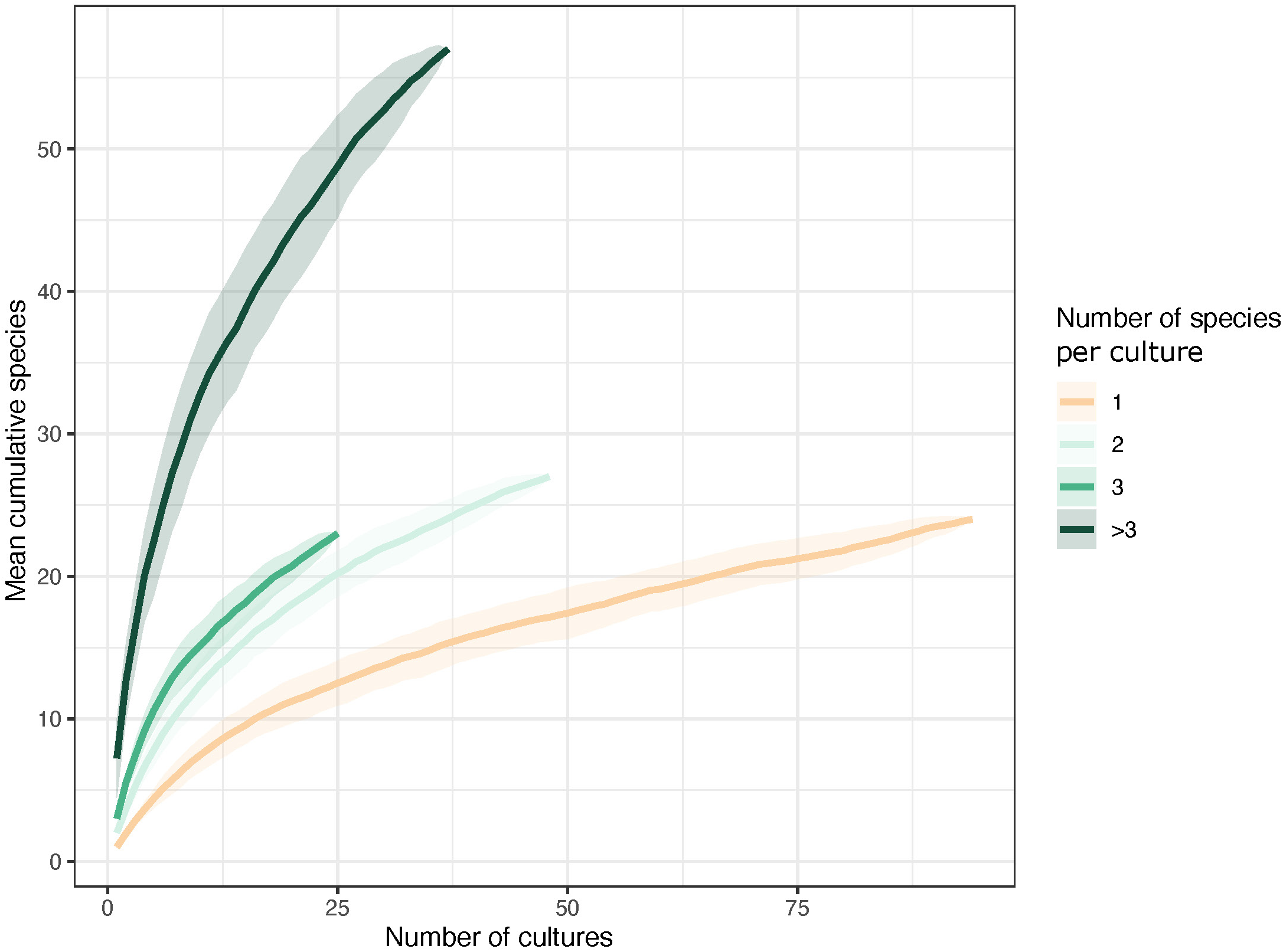
Culturing in groups enables access to a greater diversity of microbial species. Rarefaction curves show the cumulative number of unique species recovered from microbial model communities grouped by community complexity (1, 2, 3, or more than 3 species), as cultured samples are progressively added. For each category, species accumulation was calculated across 100 random permutations (bootstrapped iterations), and shaded ribbons represent ±1 standard deviation from the mean. Communities with a single species (1 genome) are shown in light orange, while increasingly complex communities (≥2 genomes) are shaded in progressively darker greens.

### Species strictly found growing in groups are more abundant and prevalent and have lower biosynthetic capacity

The total number of genomes recovered per species-cluster varied considerably (**Figure 5A**), ranging from those detected only once (e.g., sc_576 and sc_639; taxa of species-clusters can be found in **Table S4** and **Figure 3**) to those found in over 40 model communities (e.g., sc_011 and sc_014). Fifteen species (21%) were present in more than 10 model communities, collectively accounting for approximately 63% of the total MAGs (330 out of 527). We found that most of the frequently retrieved MAGs showed the flexibility of growing independently or in groups (**Figure 5A** and **B**). However, our experiment did not systematically test if species only found on their own could also grow in groups, or if species found only in groups could also grow alone. Nevertheless, when we examined the relative abundance of all cultivated species across their source environmental sample, we observed a positive correlation to the total number of genomes recovered for species found in groups (**Figure 5C**). On the other hand, the two most abundant species (Pelagibacterales sc_139 and Methylacidiphilales sc_121) were found exclusively in groups but were cultivated very few times. This might reflect that while many abundant microorganisms are easier to cultivate under our model community conditions, finding the right partners or getting the right conditions might be more challenging for others.

**Figure 5.**
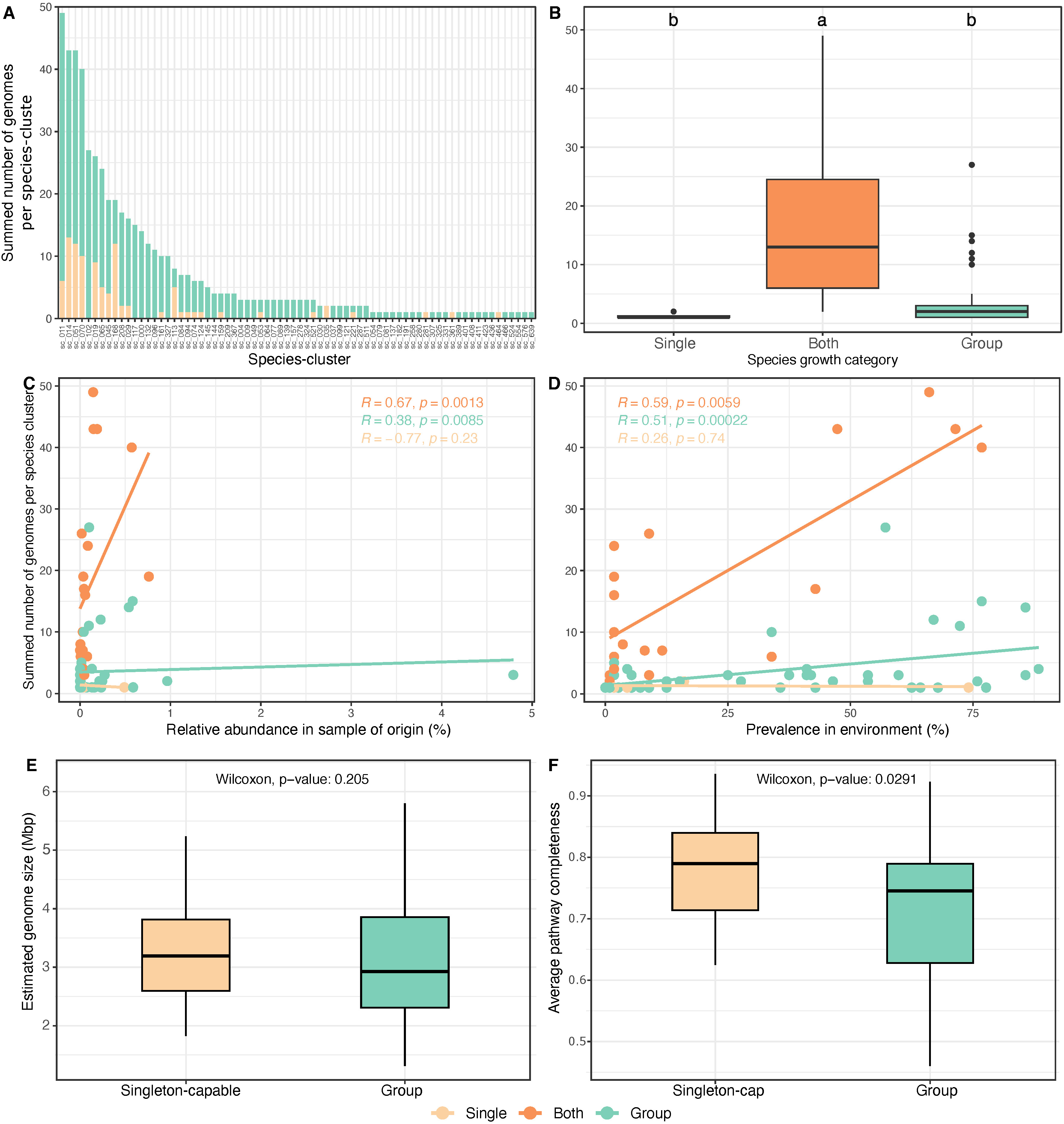
Growth strategy, environmental distribution, and genomic features of cultured microbial species. (A) Barplot showing the total number of cultured genomes recovered per species-cluster (n = 72). Bars are colored by the type of culture from which each genome was obtained: light orange for single-genome cultures and light green for multi-genome cultures. Species-clusters were further categorized based on their observed growth behavior: “Single” (light orange) for species that grew exclusively alone, “Group” (light green) for those that grew exclusively in groups, and “Both” (orange) for species that could grow alone and in groups. (B) Boxplot comparing the number of cultured genomes per species-cluster across the three growth categories (Kruskal-Wallis test, followed by Dunn’s post hoc test with Bonferroni correction; p < 0.05). Letters indicate results of Dunn’s post-hoc test (Bonferroni-corrected); groups with different letters are significantly different. (C) The dot plots display the relationship between the total number of cultured genomes per species-cluster and their relative abundance in the sample of origin (D), as well as the prevalence of these species-clusters in all environmental samples. Each data point represents one species-cluster. A non-linear regression models (Spearman) were fitted to the data. (E) Boxplot comparing the estimated genome size of species-cluster representative genomes (n = 72) growing exclusively in groups (“Group” = light green) and capable of growing alone (“Singleton-capable” = light orange). (F) Boxplot comparing the average pathway completeness of custom amino acid and vitamin biosynthesis modules between high-quality species-cluster representatives (n = 57; completeness >90%, contamination <5%) growing exclusively in groups and those capable of growing alone. Statistical significance was assessed using the Wilcoxon rank-sum test.

When examining species prevalence across all environmental samples, a similar significant positive correlation emerged with the number of cultured genomes per species (**Figure 5D**), indicating that more prevalent taxa are more frequently retrieved through our cultivation conditions.

Next, we compared estimated genome sizes and biosynthetic potential for species capable of growing alone (singleton-capable) and those that strictly grow in groups. While there was no significant difference in estimated genome sizes between these two categories (**Figure 5E**), we observed a significant difference in average pathway completeness of custom biosynthesis modules for both amino acids and vitamins, with species found strictly in groups exhibiting a lower average pathway completeness (**Figure 5F**). For this analysis, we excluded three amino acids (alanine, asparagine, and aspartate) since their biosynthesis is commonly mediated by several generic transaminases^44^ (see Methods for details). These findings suggest that the ability to grow alone is most likely associated with greater anabolic independence, whereas species that require growing in groups may depend biosynthetically on other members of the community.

### Lowest anabolic independence and higher interdependencies in species growing in groups

We next examined the biosynthetic potential of the 305 high-quality genomes (completeness >90%, contamination <5%) recovered from our microbial model communities, with a focus on how this potential relates to community complexity. We found that genomes from single-species and two-species cultures showed consistently higher anabolic independence seen in amino acid biosynthesis and vitamin modules (**Figure 6** and **Figure S8**). In contrast, genomes from three-species cultures showed lower biosynthesis potential, and cultures with more than three species have the lowest anabolic independence. This reduction was particularly pronounced in amino acid biosynthesis pathways when compared to their highest average completeness value observed in cultures with one or two species. The nine aminoacids with lowest pathway completeness were arginine (∼25% lower), proline (∼22%), phenylalanine (∼21%), tyrosine (∼20%), threonine (∼18%), leucine (∼17%), tryptophan (∼16%), serine (∼14%), and isoleucine (∼12%) (**Figure S9**). Additionally, although vitamin B12 showed a relative decrease, both single-species and more than three-species model communities had low average pathway completeness (∼22% down to ∼12%), suggesting generally limited biosynthetic capacity for B12, regardless of community complexity.

**Figure 6.**
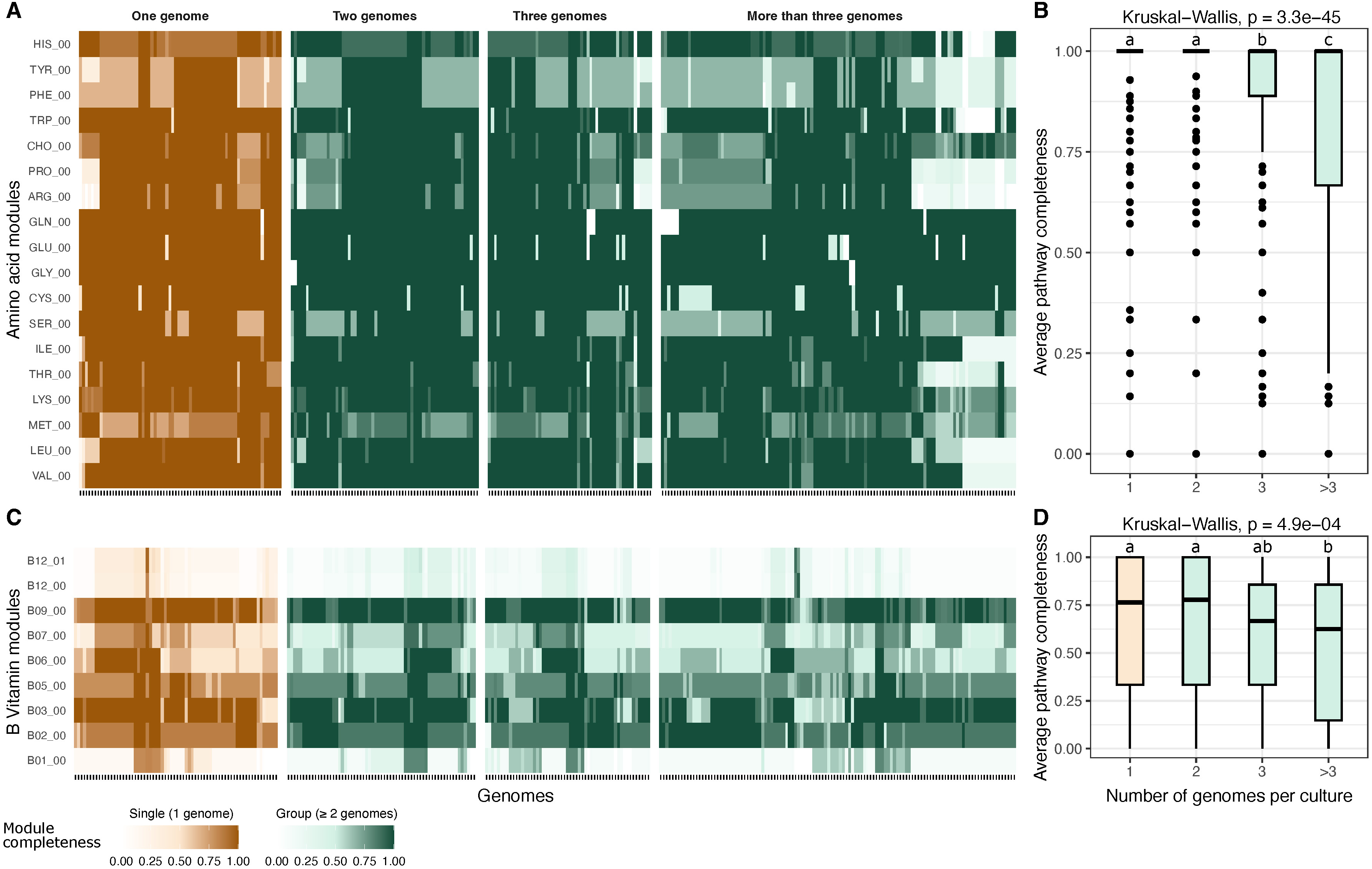
Amino acid and vitamin biosynthetic potential across species from varying community complexity. (A) Heatmap showing pathway completeness scores for 18 custom amino acid biosynthesis modules (three-letter abbreviations) across high-quality genomes (n = 305; >90% completeness, <5% contamination) grouped by community complexity: 1, 2, 3, or more than 3 genomes per culture. Each column represents a genome, and each row a biosynthetic module. Completeness values range from white (0) to dark orange (1) for genomes from single-genome cultures, and white to dark green for genomes from multi-genome cultures (≥2 genomes). (B) Boxplot summarizing the average amino acid pathway completeness per genome across the same four community complexity groups (Kruskal-Wallis test, followed by Dunn’s post hoc test with Bonferroni correction; p < 0.05). Groups with different letters are significantly different. (C) Heatmap as in panel A, but for the 9 custom vitamin biosynthesis modules. Color gradients and grouping are defined identically. (D) Boxplot summarizing the average vitamin pathway completeness per genome across complexity groups. Statistical analysis performed as in panel B.

While it is assumed that metagenome assembly might work better with lower diversity inputs, genomes from our more complex cultures (> 3 species) showed only slight differences in completeness, contamination, and N50 values (**Figure S10**). To test whether this small difference in genome quality could explain the reduced biosynthetic potential in complex communities, we examined the relationship between genome completeness and the completeness of each biosynthetic module individually (**Figure S11**). Across all 27 modules, correlations with all genomes from model communities were weak (Spearman R typically <0.3). Modules with similar completeness were often observed across genomes with varying genome completeness, indicating that minor differences in MAG quality do not drive the observed metabolic trends. We also assessed correlations for each biosynthetic module separately across the four community complexity categories (1, 2, 3, and >3 species per culture; **Figure S12**) and found only a few modules exhibited moderate correlations (R ≈ 0.3–0.5), with no systematic bias toward the more complex communities (>3 species). These results suggest that the lower biosynthetic capacity in multi-species cultures is a biological signal driven by community composition rather than a technical artifact of genome quality.

To evaluate if microbial model communities with more than three species collectively encode complete biosynthetic pathways, we evaluated gene content (based on individual module steps) of all 262 genomes recovered from 37 microbial model communities regardless of completeness. Even partial genomes were included because they can provide valuable evidence for individual metabolic steps. When examining the data at the community-level, we observed that in these cultures, all species collectively encoded the biosynthetic pathways through a mosaic of partial contributions from different species (**Figure 7**). This community-level stepwise completion indicates that biosynthetic capacity emerges collectively rather than within individual genomes, suggesting that anabolic interdependencies support the idea of facilitated community growth.

**Figure 7.**
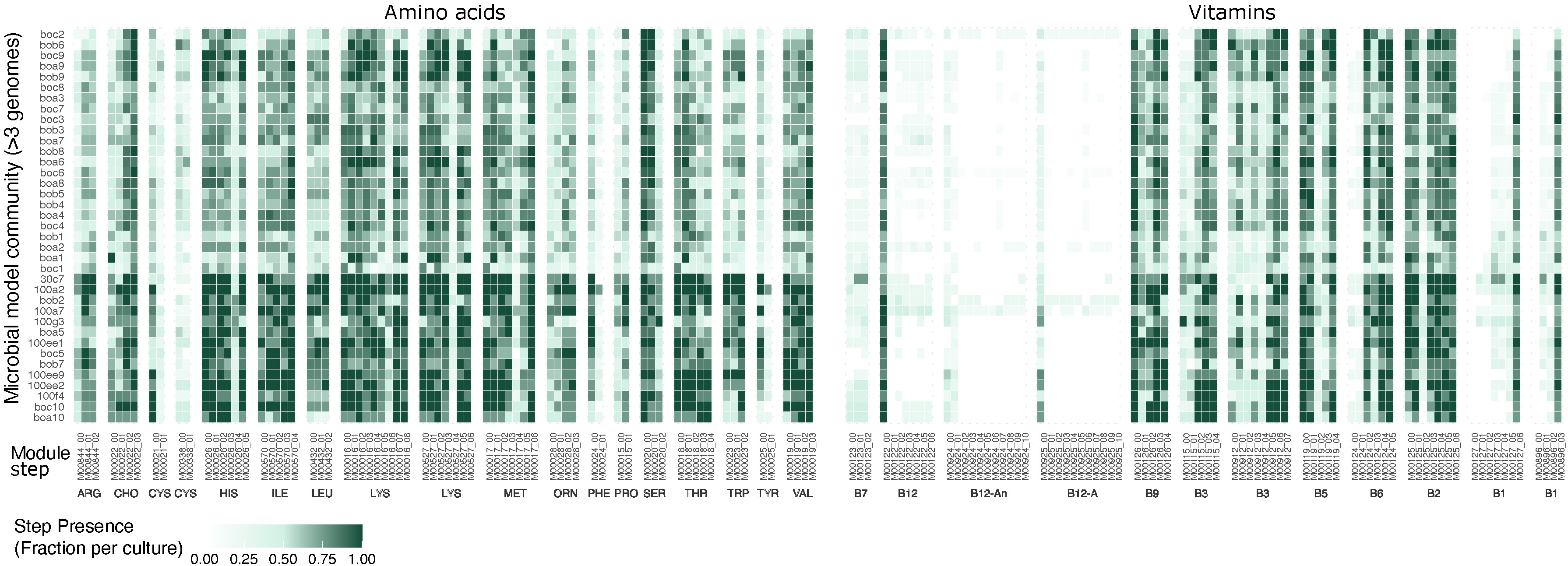
Stepwise module framework analysis of amino acid and B vitamin biosynthetic pathways across microbial model communities. Heatmap shows the average presence of biosynthetic module steps across the 204 microbial model communities, including all genomes, regardless of genome completeness. Each column represents a specific KEGG module step (e.g., M000020_01) grouped by compound. Each row corresponds to an individual model community, grouped by community complexity (1, 2, 3, or more than 3 species per culture). Values represent the proportion of genomes in each community that encode the respective module step, ranging from 0 to 1. Single-genome communities are displayed using an orange gradient (white to dark orange), while multi-genome communities (≥2 genomes) use a green gradient (white to dark green). This stepwise framework uses canonical KEGG modules (non-customized), capturing the presence of standard and alternative steps within pathways.

### Ubiquitous species tend to have smaller estimated genome sizes and reduced biosynthetic capacity

For the species from cultures (57 high-quality), we found a significant negative correlation between relative abundance and average pathway completeness for both amino acids and vitamins in species that exclusively grew in groups and those that exclusively grew alone (**Figure 8A**). This trend persisted for vitamin biosynthesis (**Figure S13C**), while amino acids alone showed a weaker association (**Figure S13B**). Moreover, we observed a significant positive correlation between estimated genome size and average pathway completeness for species that exclusively grew in groups and alone, showing that smaller genomes encode fewer biosynthetic pathways (**Figure 8B**). Notably, the correlation between vitamin biosynthesis and estimated genome size was particularly strong in group-only species (**Figure S13F**).

**Figure 8.**
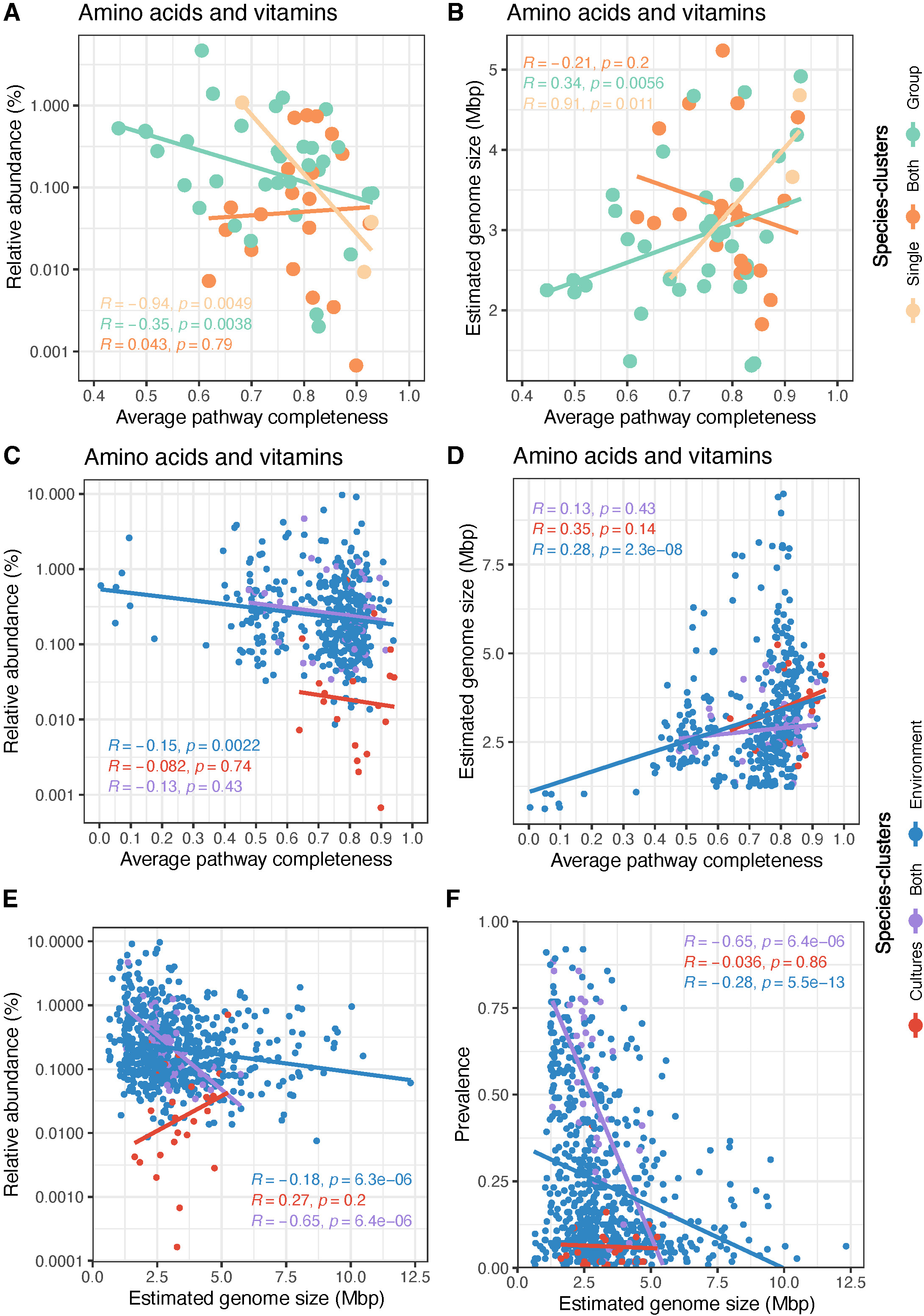
Link between pathway completeness, relative abundance, and estimated genome size of cultured species and the BalticMAG species catalog. (A) Correlation between average biosynthesis pathway completeness of custom amino acids and vitamin modules and their relative abundance of cultured species (n = 57, completeness >90%, contamination <5%). (B) Correlation between the average biosynthesis completeness for custom vitamin modules and the estimated genome size of cultured species-clusters. Each data point represents one species cluster (n = 57, completeness >90%, contamination <5%), which either grew exclusively on their own (light orange), exclusively in groups (light green), or both on their own as well as in groups (orange). Panels (C) and (D) replicate the analysis from panels (A) and (B) but include all species-clusters (n = 450, completeness >90%, contamination <5%) from both cultures and environmental samples. (E) Correlation between estimated genome size and average relative abundance and (F) correlation between estimated genome size and prevalence of all species-clusters (n = 701). Data points are color-coded to represent species clusters from cultures only (red), environments only (blue), and found in both (purple).

Scaling the observations from model communities to all species from the BalticMAG catalog (n = 450 high-quality), we found similar trends, particularly among species detected only in environmental metagenomes. In this group, we found that higher average relative abundances (**Figure 8C**) and smaller estimated genome sizes (**Figure 8D**) were linked to lower average completeness of biosynthetic pathways for amino acids and vitamins.

Finally, when we analyzed all 701 species-clusters, we observed a strong negative correlation between estimated genome size and both relative abundance (**Figure 8E**) and prevalence across samples (**Figure 8F**). These results align with prior observations, where the most successful and widespread taxa have a streamlined genome with low biosynthetic potential^17^. Collectively, these findings suggest that biosynthetic dependencies may be a common ecological strategy in the Baltic Sea.

### Anabolic dependencies in the Baltic Sea: different paths to microbial success

Given that anabolic dependencies seem to be common in the Baltic Sea, an important question arises: Are these dependencies uniform across different microorganisms, or do different microorganisms adopt distinct metabolic strategies to achieve ecological success? To explore variation in biosynthetic potential across the BalticMAG catalog, we analyzed the 450 high-quality species genomes. A principal component analysis (PCA) on the biosynthetic completeness matrix revealed a clear separation of genomes into three distinct biosynthetic completeness groups: Low (0–30%), Medium (30–62.5%), and High (>62.5%) (**Figure 9A&D**). Estimated genome size followed the trend, increasing from low to high across the biosynthetic completeness groups (**Figure 9E**). In general, among the most frequently incomplete or partially incomplete amino acid pathways were histidine, phenylalanine, and tyrosine as well as vitamin B1 and B12 (**Figure S14**). The microorganisms in the different biosynthetic groups showed distinct and clear taxonomic signatures (**Figure 9C**). The low biosynthesis completeness group consisted exclusively of *Patescibacteria* and a single *Firmicutes* genome, both lineages associated with symbiotic or highly host-dependent lifestyles. The medium biosynthesis completeness group was dominated by *Bacteroidota* and *Planctomycetota*, while the high biosynthesis group encompassed a broader diversity of phyla, including *Proteobacteria* and *Actinobacteria* as major contributors. Interestingly, microorganisms with low and high biosynthetic completeness showed no significantly different relative abundance, possibly because of the small number of taxa in the former group (**Figure 9F**).

**Figure 9.**
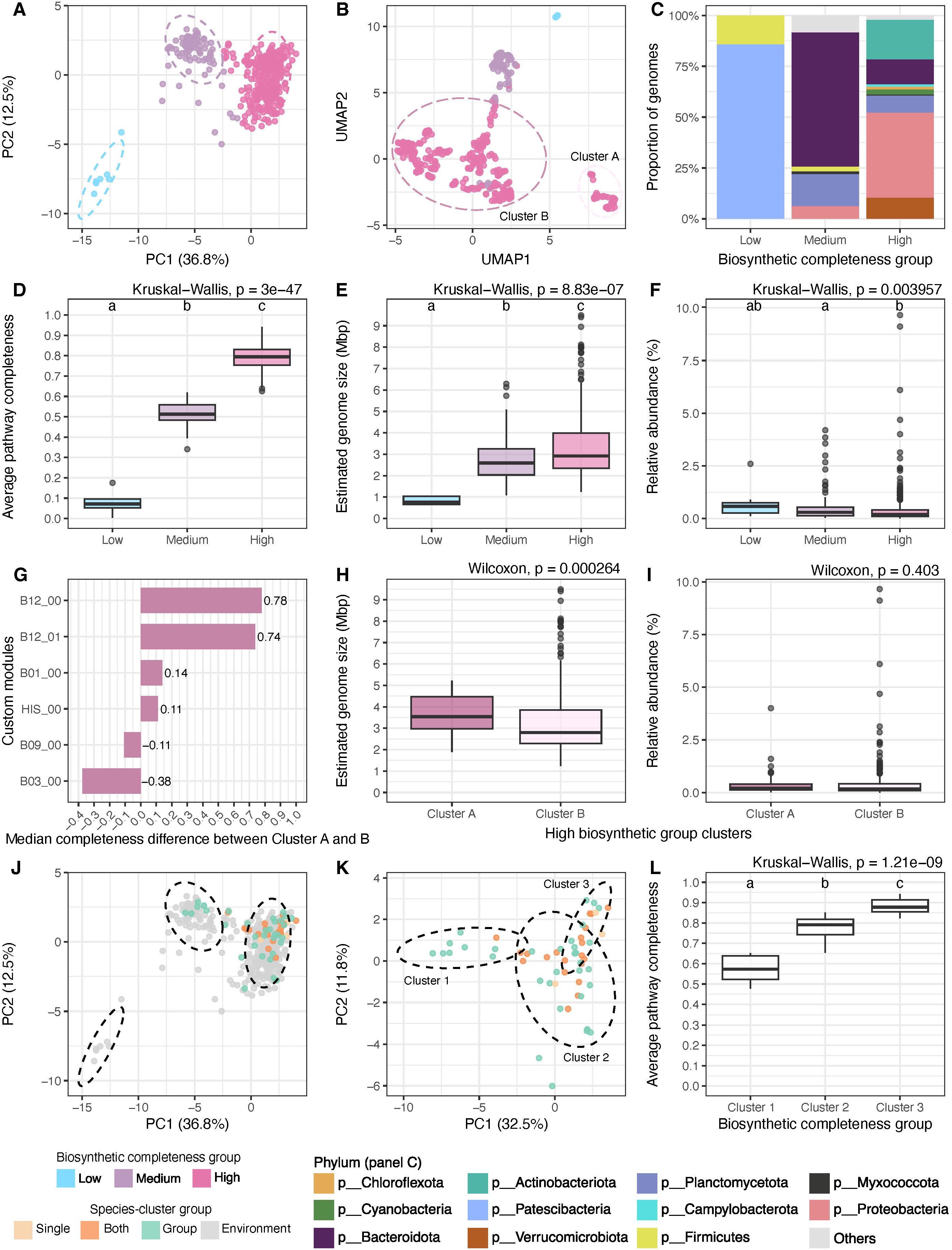
Anabolic dependencies and biosynthetic clustering of the BalticMAG species catalog inlcuding cultivated species. (A) Principal component analysis (PCA) of the biosynthetic module completeness, with genomes color-coded by biosynthetic completeness group: Low = light blue, Medium = light purple, High = pink. (B) Uniform Manifold Approximation and Projection (UMAP) of the same data, confirming the biosynthetic groupings. (C) Stacked bar plot showing the proportion and taxonomic composition of the three biosynthetic completeness groups, color-coded by phylum. (D–F) Boxplots comparing (D) average pathway completeness, (E) estimated genome size (Mbp), and (F) relative abundance of species-clusters grouped by biosynthetic completeness. Statistical significance was assessed using a Kruskal–Wallis test (p < 0.05) followed by Dunn’s post hoc test with Bonferroni correction; groups with different letters are significantly different (p < 0.05). (G) Median completeness differences for KEGG custom biosynthesis modules between Cluster A and Cluster B genomes of the High biosynthetic group. Positive values indicate modules are more complete in Cluster A, while negative values indicate modules are more complete in Cluster B. (H–I) Boxplots comparing (H) estimated genome size (Mbp) and (I) relative abundance of species-clusters from Cluster A (dark pink) and Cluster B (light pink). Statistical significance was assessed using the Wilcoxon rank-sum test (p < 0.05). (J) PCA of biosynthetic completeness color-coded by species-cluster growth category: Single = light orange, Group = light green, Both = orange; environmental-only clusters are shown in grey. (K) PCA of the 57 high-quality genomes from microbial model communities, also color-coded by growth category. (L) Boxplot showing the average pathway completeness of the three biosynthetic clusters derived from cultivated species. Statistical significance was tested using Kruskal–Wallis with Dunn’s post hoc test; groups with different letters are significantly different (p < 0.05).

To better visualize biosynthetic strategies, we projected the data using Uniform Manifold Approximation and Projection (**Figure 9B**), which further confirmed the biosynthetic groupings, with strong clustering patterns aligned with biosynthetic capacity. Interestingly, the high biosynthetic group is further separated into two subclusters. One of them, composed of 51 genomes, was taxonomically diverse yet tightly grouped. We refer to this subcluster as Cluster A, and the remaining genomes from the High biosynthetic group as Cluster B. Cluster A genomes had significantly larger genome sizes than those in Cluster B (**Figure 9H**) and appeared to be the main producers of vitamin B12 (**Figure 9G** and **Figure S15**). Cluster B genomes showed higher completeness for vitamin B3.

Finally, from the 57 high-quality genomes from cultures, we observed that we cultivated no microorganisms from the low biosynthetic group. This might suggest that cultivation techniques to grow them together with their possible host might be needed. While, most of the microorganisms we cultivated belonged to the high biosynthetic group (**Figure 9J**), the six microorganisms we cultivated from the medium biosynthetic group grew exclusively in groups. Further, a PCA based on only the genomes from the microorganisms we cultivated revealed a new set of clusters with biosynthesis values ranging from 50% to 90% (**Figure 9K** and **9L**). Cluster 1, which had the lowest biosynthesis potential, was again mostly composed of species that were capable of growing exclusively in groups. Cultivating a microorganism with lower biosynthetic potential in pure culture might indicate residual amino acids and vitamins in the filtered sterilized ocean water used as media. Nevertheless, our cultivation strategy still showed that microorganisms with lower anabolic independence prefer to grow in groups.

## DISCUSSION

The field of microbiology has traditionally relied on the isolation of microorganisms from nature in pure culture^46–51^. The practice has yielded foundational insights into microbial physiology^52–54^, metabolism^55–58^, and ecological interactions such as mutualism^28,59,60^, and competition^61–63^. However, this reductionistic approach has limitations, particularly a strong cultivation bias favoring microorganisms with larger genome sizes^41,64^, broader biosynthetic potential^65^, and greater independence. As a result, a substantial portion of the most abundant microbes in natural environments remains uncultivated^66^.

Here, we demonstrate the utility of microbial model communities established through dilution cultivation from a Baltic Sea pelagic sample for obtaining a wide range of previously uncultivated but abundant and biosynthetically limited microorganisms. Despite using only one type of ocean water as media (which did not allow us to control the presence of amino acids or vitamins), our method allowed for the cultivation of groups of microorganisms that showcased important ecological principles that govern microbial life in nature. Our results suggest that by increasingly using high-throughput dilution cultivation of microbial model communities, more diverse microorganisms with medium to high biosynthesis potential could be cultivated. Moreover, by varying cultivation physicochemical parameters, such as adding catalase^67^, changing light regimes^68^, or temperature^69^, and perhaps by leveraging a better understanding of bacterial host dynamics^70,71^, we will be able to cultivate a greater proportion of the abundant microorganisms found in nature.

Additionally, we found that species growing in microbial model communities composed of more than three species exhibited lower biosynthetic potential for both amino acids and B vitamins compared to species found growing in smaller groups or independently. The reduced per-genome biosynthetic capacity in more complex communities suggests that microorganisms with low anabolic independence are forming metabolic networks to support their nutritional requirements^72^. Interestingly, a previous study has also found a threshold of microbial diversity at which competition and complementation saturate^30^. Specifically, they observed that beyond approximately 26 taxa, further increases in diversity had little detectable impact on an overall community function (respiration), suggesting a saturation of functional capacity. Although our work examines only amino acid and vitamin biosynthesis, both studies observe a diversity threshold. These different thresholds observed emphasize that microbial communities likely achieve a balanced state of interaction complexity beyond certain diversity levels, thereby optimizing ecological efficiency.

Our findings also align with the Black Queen Hypothesis, which posits that certain functions, particularly costly biosynthetic pathways, can be lost by some community members as long as the production of the metabolite at the community level is sufficient to sustain the individual^73^. In this context, the reduced genome sizes observed in the abundant species from the model communities and the environment likely reflect a genome streamlining process^74^. The findings further align with the hypothesis that obligate co-existing microbes have evolved to rely on their community for essential nutrients^75^, potentially creating social networks that might bring microbial community stability^14,76–78^. To support this, our stepwise module analysis revealed that microbial model communities with more than three species collectively maintain biosynthetic potential for amino acids and vitamins. These observations also align with recent findings^79^ from a study which examined the potential for metabolic complementarity among auxotrophic soil bacteria. The study analyzed 746 auxotrophic strains from 27 soil-derived communities that were grown in groups of 2, 3, 4, and 5 strains, and described a clear trend: larger groups of bacteria were more capable of collectively producing all necessary amino acids due to metabolic complementarity.

Importantly, our findings also complement large-scale studies investigating the ecological distribution of auxotrophies. A recent analysis of over 26,000 representative bacterial genomes across diverse environments found that auxotrophy is more common in host-associated environments while relatively rare in aquatic and soil ecosystems. However, this analysis draws a robust but strict boundary on how to categorize microorganisms in a binary model of prototrophs and auxotrophs (more than 40% genes missing per pathway to classify as auxotroph)^16^. While this categorization has been widely used, it might also be hiding potential interdependencies. In our study, we draw no boundary for defining auxotrophy. Instead, we studied the pathways by observing their completeness and comparing them across the different species in the dataset. We believe this has the potential to reveal a wide spectrum of possibilities where microorganisms might just need a precursor to complete biosynthesis^21^.

Moreover, we found further support for the idea that metabolic interdependencies go beyond the simple exchange of end-products in a recent study^20^. The authors used 25 engineered strains of *E*. *coli* that were auxotrophic for specific amino acids (arginine, histidine, isoleucine, proline, and tryptophan) and performed pairwise cocultures of strains auxotrophic for the same amino acid. Strikingly, they found that growth complementation was often achieved by sharing the intermediates within the biosynthetic pathway. For this reason, we believe that moving away from binary categorization of auxotrophies might bring more nuance to the study of anabolic dependencies.

In summary, our study highlights the larger potential of microbial model communities in bridging the gap between laboratory cultivation and the environment. By cultivating naturally assembled groups of microorganisms, we recovered ecologically dominant taxa with limited biosynthetic capacity that are often overlooked by traditional isolation techniques. These findings demonstrate that increasing community complexity is associated with different forms of reduced metabolic autonomy. Moreover, we observed that biosynthetic interdependencies are widespread in nature. In combination with recent experimental evidence showing the exchange of biosynthetic intermediates among bacteria, our results reinforce the idea that anabolic dependencies, rather than complete autonomy, are a successful ecological strategy. Cultivating groups rather than individuals can offer a more ecologically relevant understanding of how microbes survive, interact, and evolve in nature.

## MATERIALS AND METHODS

### Sampling and sample processing

We collected an environmental sample from the surface layer of the Baltic Sea close to Askö in the Trosa archipelago (Lat 58°48.20’N, Lon 17°37.42’E) in October 2021 (**Figure 1A**). For details on the chemical composition of the water at sampling time, please visit https://shark.smhi.se/hamta-data/. The sample for our experiments was subsequently processed in the laboratory for DNA extraction and metagenomic sequencing. Briefly, the water sample was filtered through a 0.1 µm membrane and used to extract environmental DNA using either the FastDNA® SPIN Kit for Soil (MP Bio) and the Dneasy PowerWater kit (Qiagen). Additionally, we filtered water through a 0.1 µm hollow fiber cartridge (Cytiva) and used the filtrate as media to establish cultures with different starting inoculum sizes.

### Flow cytometry

We used a CytoFLEX instrument manufactured by Beckman Coulter to process the environmental sample and to calculate the cell count as the number of events/mL. Briefly, we stained 50 µl of the samples with Syto13 at a final concentration of 0.025 mM. During the flow cytometer acquisition process, we set the following parameters: FSC 2500, SSC 2500, and FITC 800 with a flow rate of 60 µl/min.

### Establishment of microbial model communities

We used the dilution-to-extinction technique considering the cell count information from the sample and diluting the cells in the filtered sterilized water, to achieve approximately the desired starting number of cells in our microbial model communities. The model communities were designed in two ways: low inoculum size and high inoculum size. The low inoculum size model communities included hundreds of individual cultures in 96-well plates with a starting number of cells of 0 (control), 2, 6, 10, 15, 20, 30, 50, and 100 cells/well. The high inoculum size model communities were prepared in bottles with starting numbers of cells of 0 (control), 200, 600, 1000, 1500, 2000, 3000, 5000, 10000, 100000, and 1000000 cells/bottle. Filtered water from the original environmental sample, without any additional nutrients, was used as media, resulting in an undefined medium closely reflecting in situ conditions. All cultures were incubated with light/dark cycles (light 6:42, dark 18:32 each day) and 12.2 °C at light and 11.8 °C at dark for four weeks before further processing. This regime was designed to emulate as best as possible the natural fluctuations in time, temperature, and light that cells experience in their environment.

### MDA, library preparation, and sequencing

Before sending our samples for metagenomic sequencing, we performed multiple displacement amplification (MDA) on all cultures to increase the concentration of DNA. The MDA reaction consisted of 0.6 µL of culture and 4.4 µL of reaction mix using the Repli-g Single Cell kit (Qiagen). After DNA amplification, we found that 315 of the cultures passed the amplification threshold of the negative controls set in the MDA reaction. These cultures were deemed positive and were sent for sequencing. We extracted DNA from the selected cultures and our environmental sample with two different DNA extraction methods and subjected them to library preparation using the TruSeq PCRfree DNA library preparation kit (Illumina Inc.) followed by metagenomic Illumina sequencing at the SNP&SEQ Platform at Uppsala University. This sequencing technology utilized cluster generation and 150 cycles of paired-end sequencing on an SP flow cell, employing the NovaSeq 6000 system with v1.5 sequencing chemistry (Illumina Inc).

### Genome resolved metagenomics pipeline

We removed low-quality reads from the raw sequences using the software Trimmomatic (v0.36)^80^ with the following options: ILLUMINACLIP:TruSeq3-PE-2.fa:2:30:10:2:keepBothReads LEADING:3 TRAILING:3 SLIDINGWINDOW:4:15 MINLEN:50. We employed the MetaWRAP pipeline (v1.3.2)^81^ to process our clean metagenomic reads. First, clean reads were assembled in a single-sample assembly style using the “metaWRAP_assembly” module with MegaHit (v1.1.3) for the environmental samples^82^ and with SPAdes (v3.15.3)^83^ for the culture samples. The quality of the assemblies was assessed with QUAST (v.5.0.2)^84^. Since the culture DNA was amplified using MDA, the read information for coverage could not be used. For this reason, we included background metagenomic data from previous projects in the Baltic Sea (**Figure 1A** and **Table S2**) for binning. Subsequently, we used multiple-sample coverage binning to decrease the contamination and increase the completeness of bins^85^. The reads were mapped against all assemblies using the Input_POGENOM pipeline^86^, which uses Bowtie2^87^ with default parameters. After mapping and obtaining the BAM files, the min coverage was calculated using samtools (v1.9)^88^. Only coverage values for each assembly in each metagenomic sample with mean coverage ≥20× and mean breadth ≥40% were retained, following Input_POGENOM recommendations. These coverage values for each sample were combined and processed with the “metaWRAP_binning” module, which uses three metagenomic binning tools: metaBAT2^89^, maxBIN2^90^, and CONCOCT^91^. We consolidated all the bins generated by these different tools using the “metaWRAP_bin_refinement” module. We classified the resulting bins taxonomically with GTDB-tk v2.1.1^92^. Finally, we assessed the quality of the bins using CheckM (v1.1.3)^93^. We considered bins as MAGs when they had a completeness of >45% and a contamination of <10%, and these MAGs were included for further analysis.

### Complementing with previously published MAGs

The de-replicated MAGs obtained here were supplemented with 771 MAGs from an earlier study^37^ that were based on metagenomics data from three studies^37–39^. We de-replicated the collection of MAGs to obtain species-cluster representatives using ANI > 95% with mOTUpan (v0.3.2)^94^, and selected the genome with the highest quality as the species-cluster representative genome.

### Relative abundance analysis

To calculate the relative abundance of our 701 species-clusters in the BalticMAG catalog, we employed the mapping tool Strobealign (v0.14.0)^95^ , which aligned the short metagenomic reads to our species-clusters collection using a high-speed indexing method (**Table S5**). Briefly, we created three different indexes with different lengths (100, 125, and 150) for our 701 species-clusters. We filtered out low-quality reads from our 112 environmental samples with Trimmonatic (v0.36)^80^ with the following options: ILLUMINACLIP:TruSeq3-PE-2.fa:2:30:10:2:keepBothReads LEADING:3 TRAILING:3 SLIDINGWINDOW:4:15 MINLEN:50. After this, we did a competitive mapping of all reads against our created index to obtain the corresponding BAM file for each sample. We sorted the BAM files with the Anvi’o platform (v7.1)^96^ using the ‘anvi-init-bam’ program, and we calculated the coverage of each genome per sample with the program ‘anvi-profile-blitz’. We calculated each species-cluster’s relative abundance with the previously obtained output by dividing each genome’s mean coverage inner quartiles (i.e., q2q3_cov) by the overall sample mean coverage. We also computed the “prevalence”, defined as the frequency of each species-cluster across samples. Specifically, prevalence represents the proportion of samples in which a species-cluster was detected with a relative abundance > 0.

### Custom functional annotation of KEGG Orthologs (KOs) and biosynthetic modules

We used the Anvi’o platform (v7.1)^97^ to perform functional annotation of KEGG Orthologs (KOs) and to estimate metabolic potential. Initially, for each genome we used ‘anvi-gen-contigs-databasè to create a contigs database, which served as the basis for the subsequent functional annotation steps. To annotate each genome with KOs from the KEGG KOfam database^98^, we ran the ‘anvi-run-kegg-kofams’ program. We then predicted the metabolic capabilities of each genome by running the ‘anvi-estimate-metabolism’ program ^99^ (**Table S6**).

In addition to default KEGG modules^100,101^, we developed and implemented a custom set of 30 curated modules targeting the biosynthesis of amino acids and B vitamins (**Table S7**, **S8**, and **S9**). These custom modules consolidate multiple KEGG modules and fill gaps for reactions without KEGG definitions. Custom module definitions and implementation files are publicly available in the accompanying GitHub repository: https://github.com/ivagljiva/custom_biosynthesis_modules.

Custom modules were integrated using the ‘anvi-setup-user-modules’ command. Completeness scores for each genome and custom module were calculated using ‘anvi-estimate-metabolism’ with the “--only-user-modules” flag. Although we initially created custom modules for all 20 proteinogenic amino acids (plus the important precursor chorismate), we subsequently excluded alanine, asparagine, and aspartate modules from downstream analysis. These three amino acids are commonly produced by generic transamination reactions with central metabolic intermediates (e.g., pyruvate or oxaloacetate), and their biosynthesis often involves multiple redundant enzymes that are still challenging to annotate accurately^44^. Because our custom definitions included only a very limited subset of these enzymes, we observed artificially low completeness scores for these three modules. Thus, for accuracy and consistency, we retained only 18 amino acid modules (plus the 9 B vitamin modules) for downstream statistical comparisons (**Table S10**).

### Statistical analyses

All statistical analyses from this study were performed with R version (v4.4.0)^102^ and RStudio^103^. We used the Shapiro test to assess the normality of our data to be compared; if p < 0.05, we interpret this as not normally distributed^104^. Since our data was not normally distributed, we employed a non-parametric test, such as the Wilcoxon test, to find differences between pairs of groups (e.g., culture vs environment)^105^. To assess statistical differences among more than two groups, we first applied a Kruskal–Wallis rank-sum test to determine if any group differed significantly^106^. When the Kruskal–Wallis test was significant (p < 0.05), we performed post-hoc pairwise comparisons using Dunn’s test with Bonferroni correction for multiple testing^107^. To visualize pairwise group differences, we applied a compact letter display (CLD), where groups that do not differ significantly share the same letter, and groups with different letters are significantly different from each other.

We also performed a Kruskal-Wallis test to compare different groups, and the p-values were corrected using the Benjamin-Hochberg procedure. Kruskal-Wallis tests were followed by Dunn’s post hoc test for pairwise comparisons, and significant differences are indicated by the different letters when p < 0.05. All comparisons were reported to be significant if the p-values were < 0.05.

## Data availability

The paired-end sequences of both environmental (n =2) and culture (n = 204) metagenomic samples from the Baltic Sea samples from this study, along with the corresponding 827 MAGs (>45% completeness and <10% contamination), have been deposited in NCBI under the BioProject ID PRJNA1134408. All other metagenomes from the Baltic Sea (n = 110) were downloaded from public repositories, and the metadata is included in **Table S2**, including their publication reference.

## ACKNOWLEDGEMENTS

This work was funded by SciLifeLab and by the Swedish Research Council VR (grant 2022-03077). We thank Jakob Walve from the Marine Laboratory for sampling the Baltic Sea on October 12, 2021 and providing the sample to us. The authors would like to acknowledge support from the Genomics infrastructure services at Science for Life Laboratory in Uppsala. The MDA was done by Claudia Bergin at the Microbial Single Cell facility, and sequencing was performed by the SNP&SEQ Technology Platform in Uppsala. The facilities are part of the National Genomics Infrastructure (NGI) Sweden and Science for Life Laboratory. The SNP&SEQ Platform is also supported by the Swedish Research Council and the Knut and Alice Wallenberg Foundation. The authors acknowledge support from SNIC/Uppsala Multidisciplinary Center for Advanced Computational Science for access to the UPPMAX computational infrastructure as well as the National Academic Infrastructure for Supercomputing in Sweden (NAISS). Computational work and data handling were enabled by resources in the projects SNIC 2022/5-392, 2023/5-126, and naiss2023-5-379 provided by the Swedish National Infrastructure for Computing (SNIC) at UPPMAX, partially funded by the Swedish Research Council through a grant agreement no. 2018-05973 and 2022-06725.

SLG conceptualized, designed, and supervised the research. FB and SLG performed the cultivation work. AFA and LFD-Z provided the Baltic Sea MAGs that contributed to the genomic catalog. AP-V, JED, LFD-Z, and SLG conducted bioinformatics analysis. AR-G supported bioinformatic analysis. IV implemented the custom modules with input from AP-V. AP-V, AT, and SLG performed the data interpretation and visualization. AP-V led the writing of the manuscript with input from SLG. All co-authors contributed to the literature searches, participated in editing and reviewing the manuscript, and approved the final version.

## REFERENCES

1. Konopka, A. What is microbial community ecology? The ISME Journal 3, 1223–1230 (2009).

2. Falkowski, P. G., Fenchel, T. & Delong, E. F. The Microbial Engines That Drive Earth’s Biogeochemical Cycles. Science 320, 1034–1039 (2008).

3. Gralka, M. Searching for Principles of Microbial Ecology Across Levels of Biological Organization. Integrative And Comparative Biology 63, 1520–1531 (2023).

4. Herman, M. A. et al. A Unifying Framework for Understanding Biological Structures and Functions Across Levels of Biological Organization. Integrative and Comparative Biology 61, 2038–2047 (2022).

5. O’Toole, G. A. We have a community problem. J Bacteriol e00073–24 (2024) doi:10.1128/jb.00073-24.

6. Mee, M. T., Collins, J. J., Church, G. M. & Wang, H. H. Syntrophic exchange in synthetic microbial communities. Proc. Natl. Acad. Sci. U.S.A. 111, (2014).

7. Mao, Z. et al. The selection of copiotrophs may complicate biodiversity-ecosystem functioning relationships in microbial dilution-to-extinction experiments. Environmental Microbiome 18, 19 (2023).

8. Benoit, G. et al. High-quality metagenome assembly from long accurate reads with metaMDBG. Nat Biotechnol 42, 1378–1383 (2024).

9. Eren, A. M. & Banfield, J. F. Modern microbiology: Embracing complexity through integration across scales. Cell 187, 5151–5170 (2024).

10. Méheust, R., Castelle, C. J., Jaffe, A. L. & Banfield, J. F. Conserved and lineage-specific hypothetical proteins may have played a central role in the rise and diversification of major archaeal groups. BMC Biol 20, 154 (2022).

11. Hug, L. A. et al. A new view of the tree of life. Nat Microbiol 1, 16048 (2016).

12. Anantharaman, K. et al. Thousands of microbial genomes shed light on interconnected biogeochemical processes in an aquifer system. Nat Commun 7, 13219 (2016).

13. Davis, B. D. & Mingioli, E. S. Mutants of Escherichia coli requiring methionine or vitamin B12. J Bacteriol 60, 17–28 (1950).

14. Giordano, N. et al. Genome-scale community modelling reveals conserved metabolic cross-feedings in epipelagic bacterioplankton communities. Nat Commun 15, 2721 (2024).

15. Hessler, T. et al. Vitamin interdependencies predicted by metagenomics-informed network analyses and validated in microbial community microcosms. Nat Commun 14, (2023).

16. Ramoneda, J., Jensen, T. B. N., Price, M. N., Casamayor, E. O. & Fierer, N. Taxonomic and environmental distribution of bacterial amino acid auxotrophies. Nat Commun 14, (2023).

17. Rodríguez-Gijón, A. et al. The ecological success of freshwater microorganisms is mediated by streamlining and biotic interactions. Preprint at 10.1101/2025.03.24.644981 (2025).

18. Garcia, S. L. et al. Auxotrophy and intrapopulation complementary in the ‘interactome’ of a cultivated freshwater model community. Mol Ecol 24, 4449–4459 (2015).

19. Gómez Consarnau, L., et al. Mosaic patterns of B vitamin synthesis and utilization in a natural marine microbial community. Environmental Microbiology 20, 2809–2823 (2018).

20. Hong, Y.-J., Cai, Y. & Antoniewicz, M. R. Cross-feeding of amino acid pathway intermediates is common in co-cultures of auxotrophic Escherichia coli. Metabolic Engineering 88, 172–179 (2025).

21. Wienhausen, G. et al. Ligand cross-feeding resolves bacterial vitamin B12 auxotrophies. Nature (2024) doi:10.1038/s41586-024-07396-y.

22. Oña, L. & Kost, C. Cooperation increases robustness to ecological disturbance in microbial cross feeding networks. Ecology Letters 25, 1410–1420 (2022).

23. Aziz, F. A. A. et al. Interspecies interactions are an integral determinant of microbial community dynamics. Front. Microbiol. 6, (2015).

24. Goldford, J. E., et al. Emergent simplicity in microbial community assembly. (2018).

25. Pacheco, A. R., Osborne, M. L. & Segrè, D. Non-additive microbial community responses to environmental complexity. Nat Commun 12, 2365 (2021).

26. Bayer, B. et al. Metabolite release by nitrifiers facilitates metabolic interactions in the ocean. The ISME Journal wrae172 (2024) doi:10.1093/ismejo/wrae172.

27. Grant, M. A. A., Kazamia, E., Cicuta, P. & Smith, A. G. Direct exchange of vitamin B12 is demonstrated by modelling the growth dynamics of algal–bacterial cocultures. The ISME Journal 8, 1418–1427 (2014).

28. Hillesland, K. L. & Stahl, D. A. Rapid evolution of stability and productivity at the origin of a microbial mutualism. Proc. Natl. Acad. Sci. U.S.A. 107, 2124–2129 (2010).

29. Chang, C.-Y., Bajić, D., Vila, J. C. C., Estrela, S. & Sanchez, A. Emergent coexistence in multispecies microbial communities. Science 381, 343–348 (2023).

30. Yu, X., Polz, M. F. & Alm, E. J. Interactions in self-assembled microbial communities saturate with diversity. The ISME Journal 13, 1602–1617 (2019).

31. Hammarlund, S. P., Gedeon, T., Carlson, R. P. & Harcombe, W. R. Limitation by a shared mutualist promotes coexistence of multiple competing partners. Nat Commun 12, 619 (2021).

32. Pande, S. et al. Fitness and stability of obligate cross-feeding interactions that emerge upon gene loss in bacteria. ISME J 8, 953–962 (2014).

33. Garcia, S. L. et al. Model Communities Hint at Promiscuous Metabolic Linkages between Ubiquitous Free-Living Freshwater Bacteria. mSphere 3, e00202–18 (2018).

34. Ratzke, C., Barrere, J. & Gore, J. Strength of species interactions determines biodiversity and stability in microbial communities. Nat Ecol Evol 4, 376–383 (2020).

35. Steen, A. D. et al. High proportions of bacteria and archaea across most biomes remain uncultured. ISME J 13, 3126–3130 (2019).

36. Garcia, S. Mixed cultures as model communities: hunting for ubiquitous microorganisms, their partners, and interactions. Aquat. Microb. Ecol. 77, 79–85 (2016).

37. Alneberg, J., et al. Ecosystem-wide metagenomic binning enables prediction of ecological niches from genomes. Commun Biol 3, (2020).

38. Alneberg, J. et al. BARM and BalticMicrobeDB, a reference metagenome and interface to meta-omic data for the Baltic Sea. Sci Data 5, (2018).

39. Larsson, J. et al. Picocyanobacteria containing a novel pigment gene cluster dominate the brackish water Baltic Sea. The ISME Journal 8, 1892–1903 (2014).

40. Pacheco-Valenciana, A., Garcia, S. L., Dharamshi, J. E., Delgado-Zambrano, L. F. & Andersson, A. F. The BalticMAG catalog. 10.17044/scilifelab.28746086.v1 (2025).

41. Rodríguez-Gijón, A. et al. A Genomic Perspective Across Earth’s Microbiomes Reveals That Genome Size in Archaea and Bacteria Is Linked to Ecosystem Type and Trophic Strategy. Front. Microbiol. 12, 761869 (2022).

42. Herlemann, D. P. et al. Transitions in bacterial communities along the 2000 km salinity gradient of the Baltic Sea. ISME J 5, 1571–1579 (2011).

43. Rodríguez-Gijón, A. et al. Linking prokaryotic genome size variation to metabolic potential and environment. ISME COMMUN. 3, 25 (2023).

44. Price, M. N., Deutschbauer, A. M. & Arkin, A. P. GapMind: Automated Annotation of Amino Acid Biosynthesis. mSystems 5, 10.1128/msystems.00291-20 (2020).

45. Somerville, V. et al. Long-read based de novo assembly of low-complexity metagenome samples results in finished genomes and reveals insights into strain diversity and an active phage system. BMC Microbiol 19, 143 (2019).

46. Gich, F., Schubert, K., Bruns, A., Hoffelner, H. & Overmann, J. Specific Detection, Isolation, and Characterization of Selected, Previously Uncultured Members of the Freshwater Bacterioplankton Community. Appl Environ Microbiol 71, 5908–5919 (2005).

47. Janssen, P. H., Yates, P. S., Grinton, B. E., Taylor, P. M. & Sait, M. Improved Culturability of Soil Bacteria and Isolation in Pure Culture of Novel Members of the Divisions *Acidobacteria* , *Actinobacteria* , *Proteobacteria* , and *Verrucomicrobia*. Appl Environ Microbiol 68, 2391–2396 (2002).

48. Kaeberlein, T., Lewis, K. & Epstein, S. S. Isolating ‘Uncultivable’ Microorganisms in Pure Culture in a Simulated Natural Environment. Science 296, 1127–1129 (2002).

49. Könneke, M. et al. Isolation of an autotrophic ammonia-oxidizing marine archaeon. Nature 437, 543–546 (2005).

50. Rappé, M. S., Connon, S. A., Vergin, K. L. & Giovannoni, S. J. Cultivation of the ubiquitous SAR11 marine bacterioplankton clade. Nature 418, 630–633 (2002).

51. Schut, F. et al. Isolation of Typical Marine Bacteria by Dilution Culture: Growth, Maintenance, and Characteristics of Isolates under Laboratory Conditions. Appl Environ Microbiol 59, 2150–2160 (1993).

52. Mella-Flores, D. et al. Prochlorococcus and Synechococcus have Evolved Different Adaptive Mechanisms to Cope with Light and UV Stress. Front. Microbio. 3, (2012).

53. Orr, P. T. & Jones, G. J. Relationship between microcystin production and cell division rates in nitrogen limited Microcystis aeruginosa cultures. Limnology & Oceanography 43, 1604–1614 (1998).

54. Sohm, J. A., Edwards, B. R., Wilson, B. G. & Webb, E. A. Constitutive Extracellular Polysaccharide (EPS) Production by Specific Isolates of Crocosphaera watsonii. Front. Microbio. 2, (2011).

55. Fischer, F., Zillig, W., Stetter, K. O. & Schreiber, G. Chemolithoautotrophic metabolism of anaerobic extremely thermophilic archaebacteria. Nature 301, 511–513 (1983).

56. Leschine, S. B., Holwell, K. & Canale-Parola, E. Nitrogen Fixation by Anaerobic Cellulolytic Bacteria. Science 242, 1157–1159 (1988).

57. Zehnder, A. J. & Brock, T. D. Methane formation and methane oxidation by methanogenic bacteria. J Bacteriol 137, 420–432 (1979).

58. Zhang, T., Shi, X.-C., Ding, R., Xu, K. & Tremblay, P.-L. The hidden chemolithoautotrophic metabolism of *Geobacter sulfurreducens* uncovered by adaptation to formate. The ISME Journal 14, 2078–2089 (2020).

59. Belzer, C. et al. Microbial Metabolic Networks at the Mucus Layer Lead to Diet-Independent Butyrate and Vitamin B_12_ Production by Intestinal Symbionts. mBio 8, e00770–17 (2017).

60. Kazamia, E. et al. Mutualistic interactions between vitamin B_12_ dependent algae and heterotrophic bacteria exhibit regulation. Environmental Microbiology 14, 1466–1476 (2012).

61. Chodkowski, J. L. & Shade, A. Bioactive exometabolites drive maintenance competition in simple bacterial communities. mSystems e00064–24 (2024) doi:10.1128/msystems.00064-24.

62. Harrison, F., Paul, J., Massey, R. C. & Buckling, A. Interspecific competition and siderophore-mediated cooperation in *Pseudomonas aeruginosa*. The ISME Journal 2, 49–55 (2008).

63. Rao, D., Webb, J. S. & Kjelleberg, S. Competitive Interactions in Mixed-Species Biofilms Containing the Marine Bacterium *Pseudoalteromonas tunicata*. Appl Environ Microbiol 71, 1729–1736 (2005).

64. Han, K. et al. Extraordinary expansion of a Sorangium cellulosum genome from an alkaline milieu. Sci Rep 3, 2101 (2013).

65. Bentkowski, P., Van Oosterhout, C. & Mock, T. A Model of Genome Size Evolution for Prokaryotes in Stable and Fluctuating Environments. Genome Biol Evol 7, 2344–2351 (2015).

66. Lloyd, K. G., Steen, A. D., Ladau, J., Yin, J. & Crosby, L. Phylogenetically Novel Uncultured Microbial Cells Dominate Earth Microbiomes. mSystems 3, e00055–18 (2018).

67. Kim, S., Kang, I., Seo, J.-H. & Cho, J.-C. Culturing the ubiquitous freshwater actinobacterial acI lineage by supplying a biochemical ‘helper’ catalase. ISME J 13, 2252–2263 (2019).

68. Bialevich, V., Zachleder, V. & Bišová, K. The Effect of Variable Light Source and Light Intensity on the Growth of Three Algal Species. Cells 11, 1293 (2022).

69. Jiang, L. & Morin, P. J. Temperature fluctuation facilitates coexistence of competing species in experimental microbial communities. Journal of Animal Ecology 76, 660–668 (2007).

70. Kuroda, K. et al. Microscopic and metatranscriptomic analyses revealed unique cross-domain parasitism between phylum *Candidatus* Patescibacteria/candidate phyla radiation and methanogenic archaea in anaerobic ecosystems. mBio 15, e03102–23 (2024).

71. Man, D. K. W. et al. Enrichment of different taxa of the enigmatic candidate phyla radiation bacteria using a novel picolitre droplet technique. ISME Communications 4, ycae080 (2024).

72. Johnson, W. M. et al. Auxotrophic interactions: a stabilizing attribute of aquatic microbial communities? FEMS Microbiology Ecology 96, fiaa115 (2020).

73. Morris, J. J., Lenski, R. E. & Zinser, E. R. The Black Queen Hypothesis: Evolution of Dependencies through Adaptive Gene Loss. mBio 3, e00036–12 (2012).

74. Giovannoni, S. J., Cameron Thrash, J. & Temperton, B. Implications of streamlining theory for microbial ecology. ISME J 8, 1553–1565 (2014).

75. Sokolovskaya, O. M., Shelton, A. N. & Taga, M. E. Sharing vitamins: Cobamides unveil microbial interactions. Science 369, (2020).

76. Kost, C., Patil, K. R., Friedman, J., Garcia, S. L. & Ralser, M. Metabolic exchanges are ubiquitous in natural microbial communities. Nat Microbiol 8, 2244–2252 (2023).

77. Zelezniak, A. et al. Metabolic dependencies drive species co-occurrence in diverse microbial communities. Proc. Natl. Acad. Sci. U.S.A. 112, 6449–6454 (2015).

78. Zengler, K. & Zaramela, L. S. The social network of microorganisms — how auxotrophies shape complex communities. Nat Rev Microbiol 16, 383–390 (2018).

79. Yousif, G. et al. Obligate cross-feeding of metabolites is common in soil microbial communities. Preprint at 10.1101/2025.01.29.635426 (2025).

80. Bolger, A. M., Lohse, M. & Usadel, B. Trimmomatic: a flexible trimmer for Illumina sequence data. Bioinformatics 30, 2114–2120 (2014).

81. Uritskiy, G. V., DiRuggiero, J. & Taylor, J. MetaWRAP—a flexible pipeline for genome-resolved metagenomic data analysis. Microbiome 6, 158 (2018).

82. Li, D., Liu, C.-M., Luo, R., Sadakane, K. & Lam, T.-W. MEGAHIT: an ultra-fast single-node solution for large and complex metagenomics assembly via succinct *de Bruijn* graph. Bioinformatics 31, 1674–1676 (2015).

83. Bankevich, A. et al. SPAdes: A New Genome Assembly Algorithm and Its Applications to Single-Cell Sequencing. Journal of Computational Biology 19, 455–477 (2012).

84. Gurevich, A., Saveliev, V., Vyahhi, N. & Tesler, G. QUAST: quality assessment tool for genome assemblies. Bioinformatics 29, 1072–1075 (2013).

85. Mattock, J. & Watson, M. A comparison of single-coverage and multi-coverage metagenomic binning reveals extensive hidden contamination. Nat Methods 20, 1170– 1173 (2023).

86. Sjöqvist, C., Delgado, L. F., Alneberg, J. & Andersson, A. F. Ecologically coherent population structure of uncultivated bacterioplankton. The ISME Journal 15, 3034–3049 (2021).

87. Langmead, B. & Salzberg, S. L. Fast gapped-read alignment with Bowtie 2. Nat Methods 9, 357–359 (2012).

88. Li, H. et al. The Sequence Alignment/Map format and SAMtools. Bioinformatics 25, 2078–2079 (2009).

89. Kang, D. D. et al. MetaBAT 2: an adaptive binning algorithm for robust and efficient genome reconstruction from metagenome assemblies. PeerJ 7, e7359 (2019).

90. Wu, Y.-W., Simmons, B. A. & Singer, S. W. MaxBin 2.0: an automated binning algorithm to recover genomes from multiple metagenomic datasets. Bioinformatics 32, 605–607 (2016).

91. Alneberg, J. et al. Binning metagenomic contigs by coverage and composition. Nat Methods 11, 1144–1146 (2014).

92. Chaumeil, P.-A., Mussig, A. J., Hugenholtz, P. & Parks, D. H. GTDB-Tk: a toolkit to classify genomes with the Genome Taxonomy Database. Bioinformatics 36, 1925–1927 (2020).

93. Parks, D. H., Imelfort, M., Skennerton, C. T., Hugenholtz, P. & Tyson, G. W. CheckM: assessing the quality of microbial genomes recovered from isolates, single cells, and metagenomes. Genome Res. 25, 1043–1055 (2015).

94. Buck, M., Mehrshad, M. & Bertilsson, S. mOTUpan: a robust Bayesian approach to leverage metagenome-assembled genomes for core-genome estimation. NAR Genomics and Bioinformatics 4, lqac060 (2022).

95. Sahlin, K. Strobealign: flexible seed size enables ultra-fast and accurate read alignment. Genome Biol 23, 260 (2022).

96. Eren, A. M. et al. Anvi’o: an advanced analysis and visualization platform for ‘omics data. PeerJ 3, e1319 (2015).

97. Eren, A. M. et al. Community-led, integrated, reproducible multi-omics with anvi’o. Nat Microbiol 6, 3–6 (2020).

98. Aramaki, T. et al. KofamKOALA: KEGG Ortholog assignment based on profile HMM and adaptive score threshold. Bioinformatics 36, 2251–2252 (2020).

99. Veseli, I. et al. Microbes with higher metabolic independence are enriched in human gut microbiomes under stress. eLife 12, RP89862 (2025).

100. Kanehisa, M. et al. Data, information, knowledge and principle: back to metabolism in KEGG. Nucl. Acids Res. 42, D199–D205 (2014).

101. Kanehisa, M., Furumichi, M., Tanabe, M., Sato, Y. & Morishima, K. KEGG: new perspectives on genomes, pathways, diseases and drugs. Nucleic Acids Res 45, D353– D361 (2017).

102. R Core Team. R: A Language and Environment for Statistical Computing. R Foundation for Statistical Computing (2024).

103. RStudio Team. RStudio: Integrated Development for R. (2024).

104. Royston, J. P. An Extension of Shapiro and Wilk’s W Test for Normality to Large Samples. Applied Statistics 31, 115 (1982).

105. Bauer, D. F. Constructing Confidence Sets Using Rank Statistics. Journal of the American Statistical Association 67, 687–690 (1972).

106. Kruskal, W. H. & Wallis, W. A. Use of Ranks in One-Criterion Variance Analysis. Journal of the American Statistical Association 47, 583–621 (1952).

107. Dunn, O. J. Multiple Comparisons Using Rank Sums. Technometrics 6, 241–252 (1964).

