## Supplemental figures for "Low anabolic independence emerges when cultivating more than three bacterial species together"

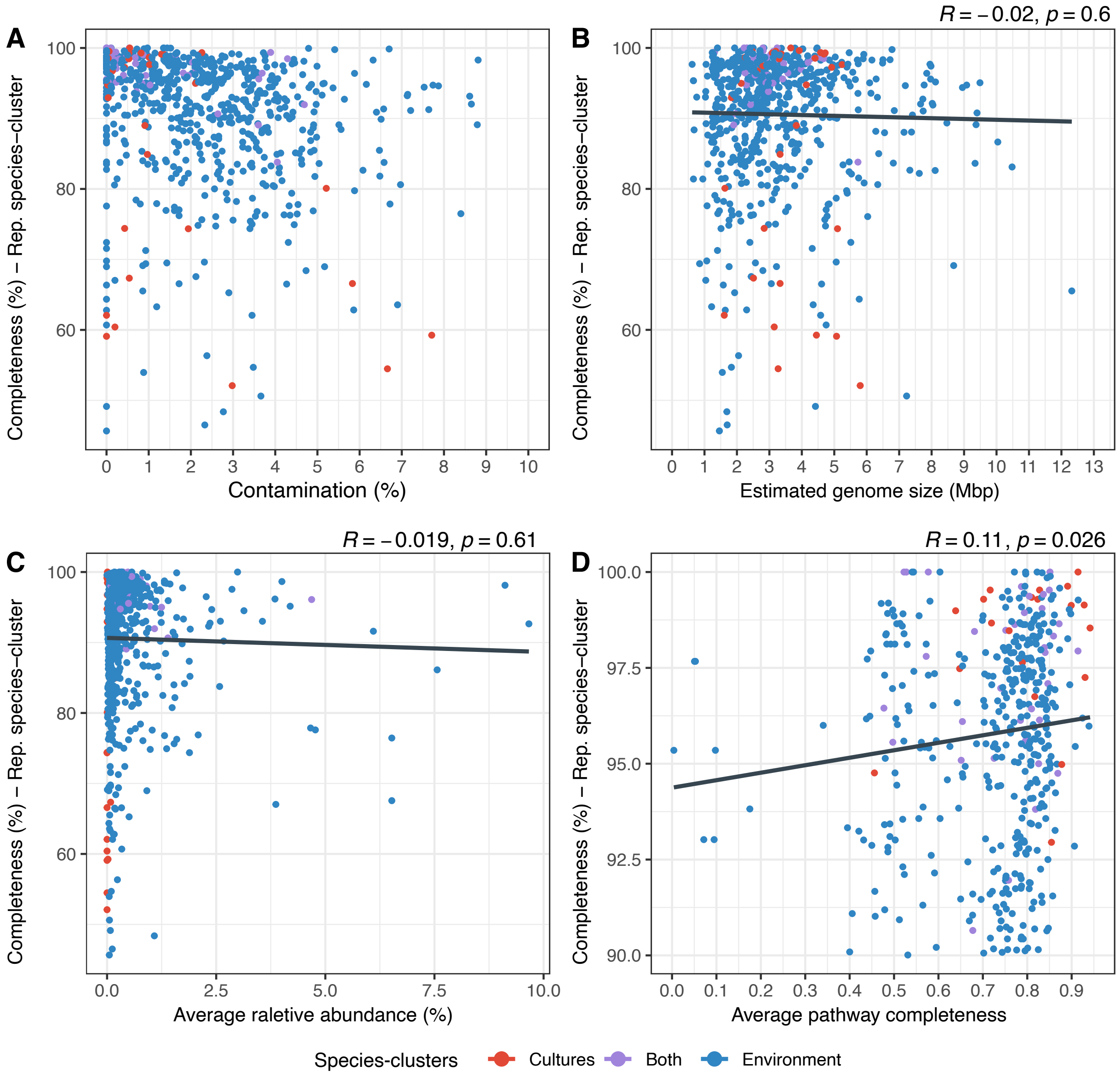


**Figure S1.** Dot plots showing the relationship between genome completeness of the BalticMAG species-clusters representatives and various metrics. These metrics include genome contamination (A), estimated genome size (B), average relative abundance (C), and average pathway completeness for both amino acids and vitamins (D). For panel D, genomes with <90% completeness and >5% contamination were filtered out for correlations using the custom average pathway completeness data. Each data point represents one species-cluster representative genome found exclusively in model communities (red), exclusively in the environment (blue), or in both (purple). Linear regression models were fitted to the data.

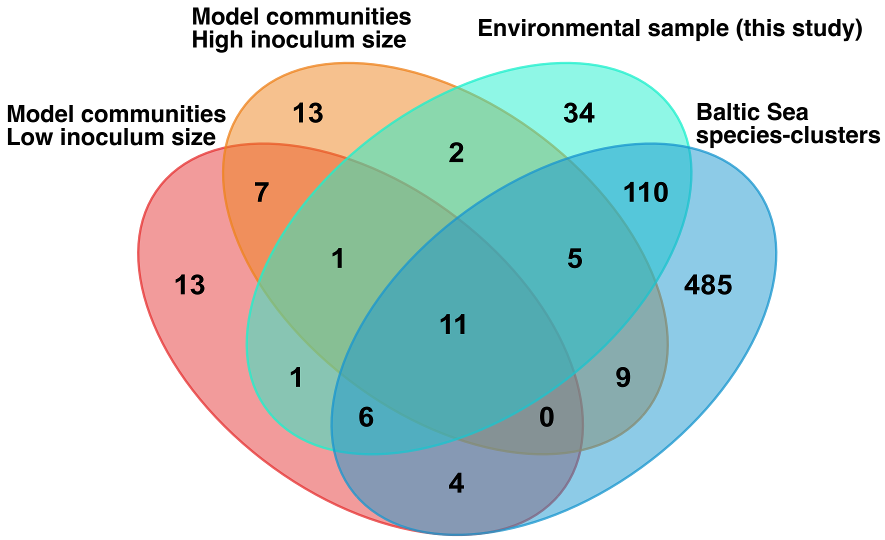


**Figure S2.** Venn diagram illustrating the overlap of species-clusters from different origins. Model communities are represented by low inoculum size (red) and high inoculum size (orange). Baltic Sea environmental samples are depicted with light blue (from this study) and dark blue (from publicly available sources). The diagram highlights the extent of overlap between these groups.

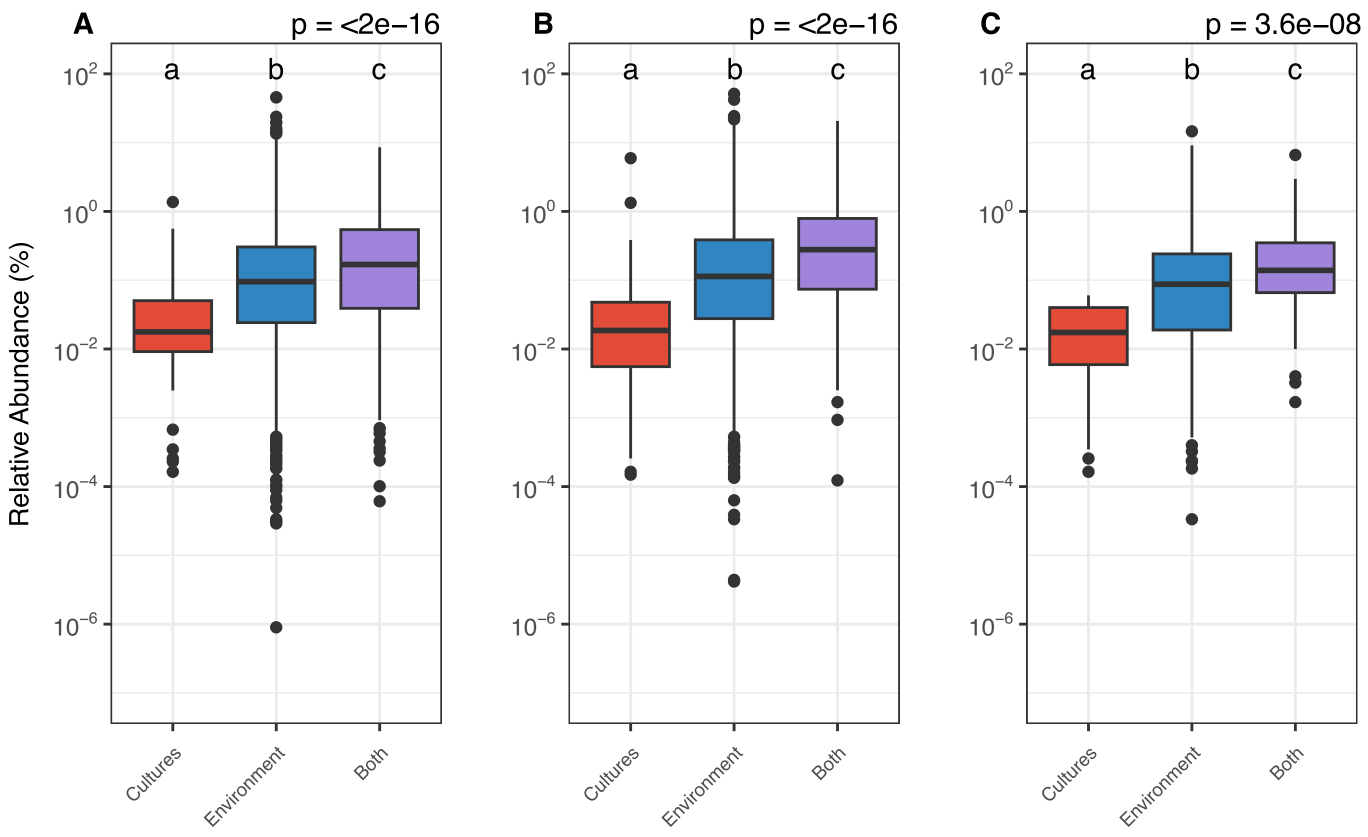


**Figure S3.** Boxplots comparing the relative abundance of species-clusters categorized by their presence in cultures (red), the environment (blue), or both (purple). (A) Relative abundances of all species across all samples with salinity between 7 and 8‰ (PSU) and (B) across only the samples from the same location as those used to establish model communities. (C) Relative abundances of species only detected in our own metagenomic samples (566 species), compared across all metagenomics samples. We used the Kruskal-Wallis test to look for statistical significance, followed by Dunn's post hoc test for pairwise comparisons. Groups with different letters are significantly different ($p < 0.05$).

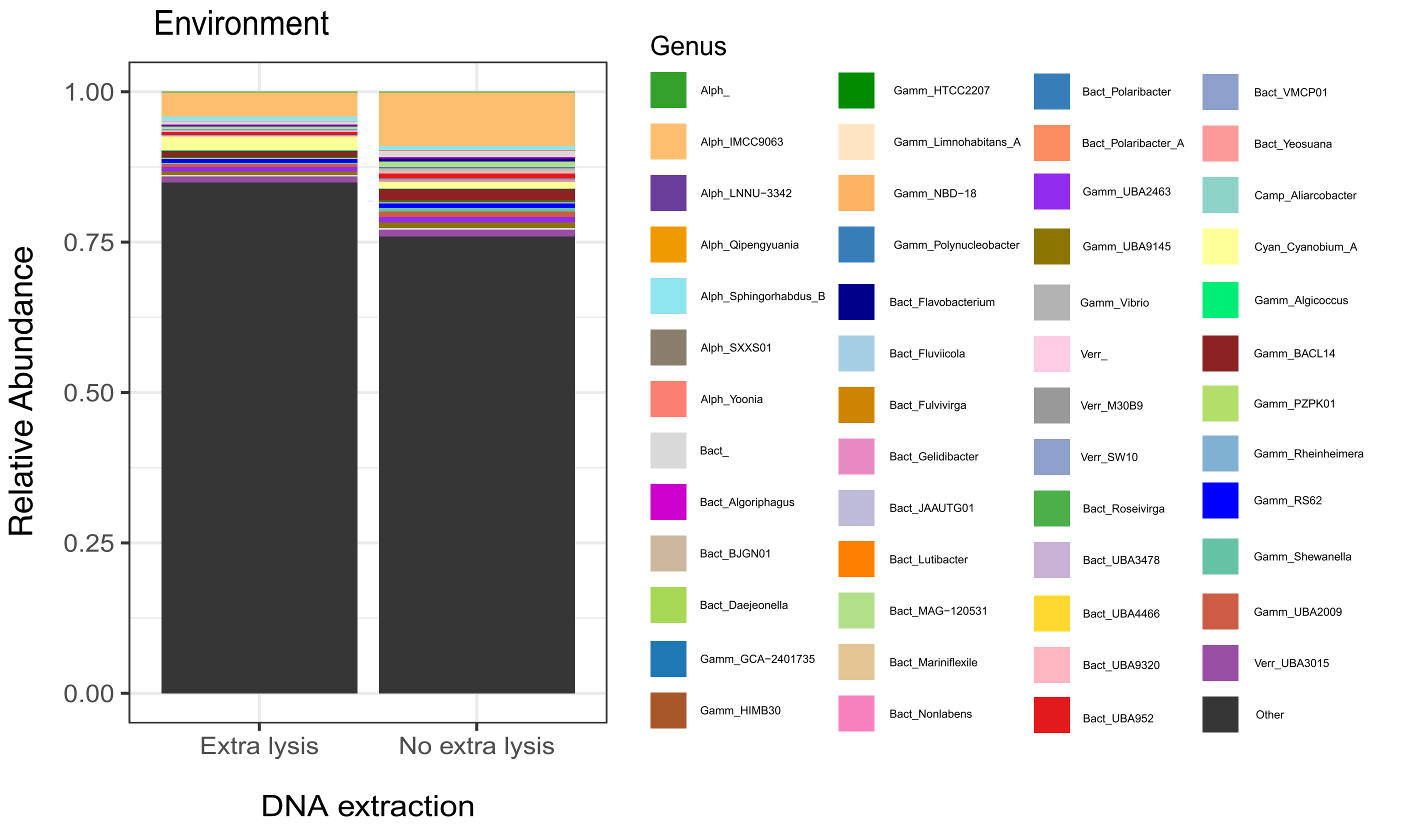


**Figure S4.** Barplot illustrating the relative abundance of environmental (dark grey) and cultivated (other colors) genomes in the sample of origin of the cultures. Note that the sample was extracted with two different DNA extraction methods, leading to two different perspectives on the diversity. Genomes are color-coded by genera, and the legend shows abbreviated prefixed indicating the class of each genus (e.g., Alph_ for Alphaproteobacteria, Bact_ for Bacteroidia, Camp_ for Campylobacteria, Cyan_ for Cyanobacteria, Gamm_ for Gammaproteobacteria, and Verr_ for Verrucomicrobiae.

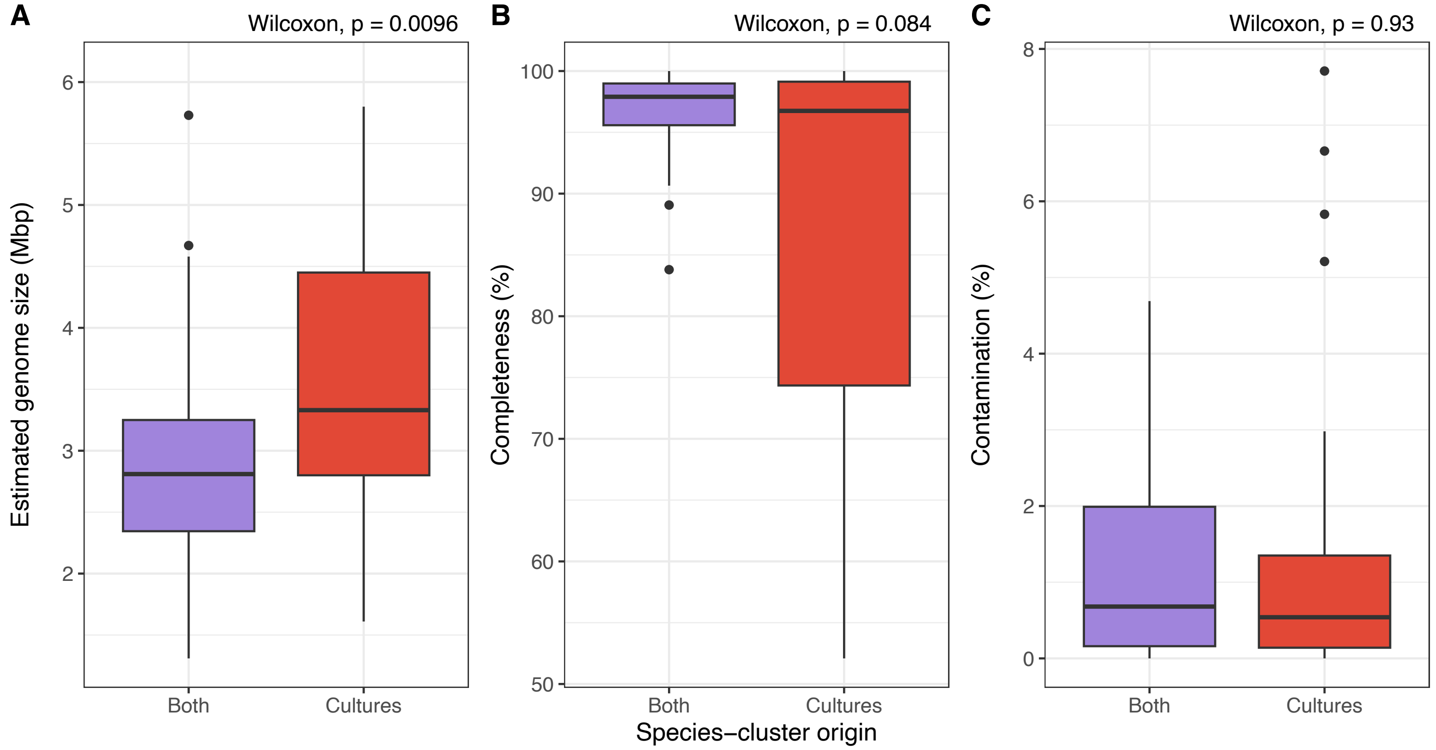


**Figure S5.** Boxplot comparing the (A) estimated genome size (Mbp), (B) completeness, and (C) contamination of species-clusters found only in cultures (red), and in both cultures and the environment (purple). Statistical significance was tested using the Wilcoxon rank-sum test ($p < 0.05$).

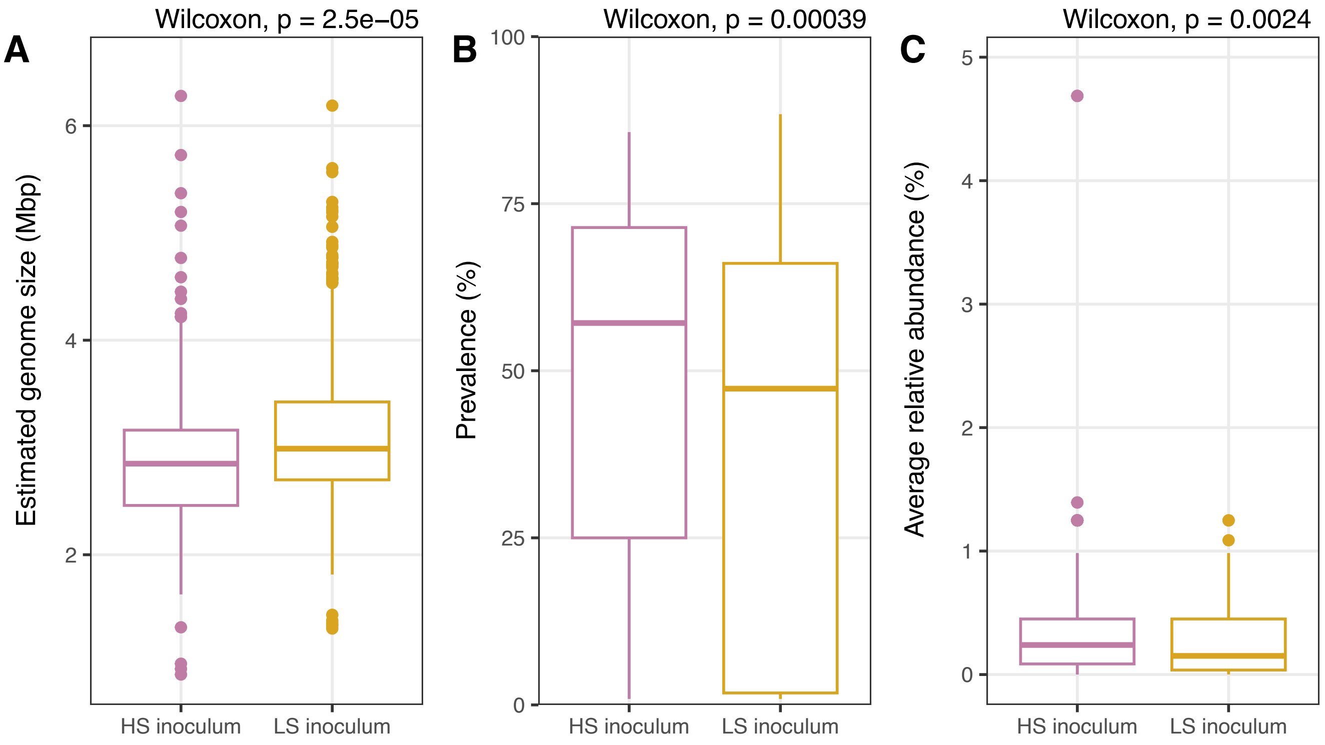


**Figure S6.** Boxplots comparing the (A) estimated genome size, (B) prevalence, and (C) average relative abundance of genomes from model communities with a high inoculum size (HS) and a low inoculum size (LS). Statistical significance was tested using the Wilcoxon rank-sum test ($p < 0.05$).

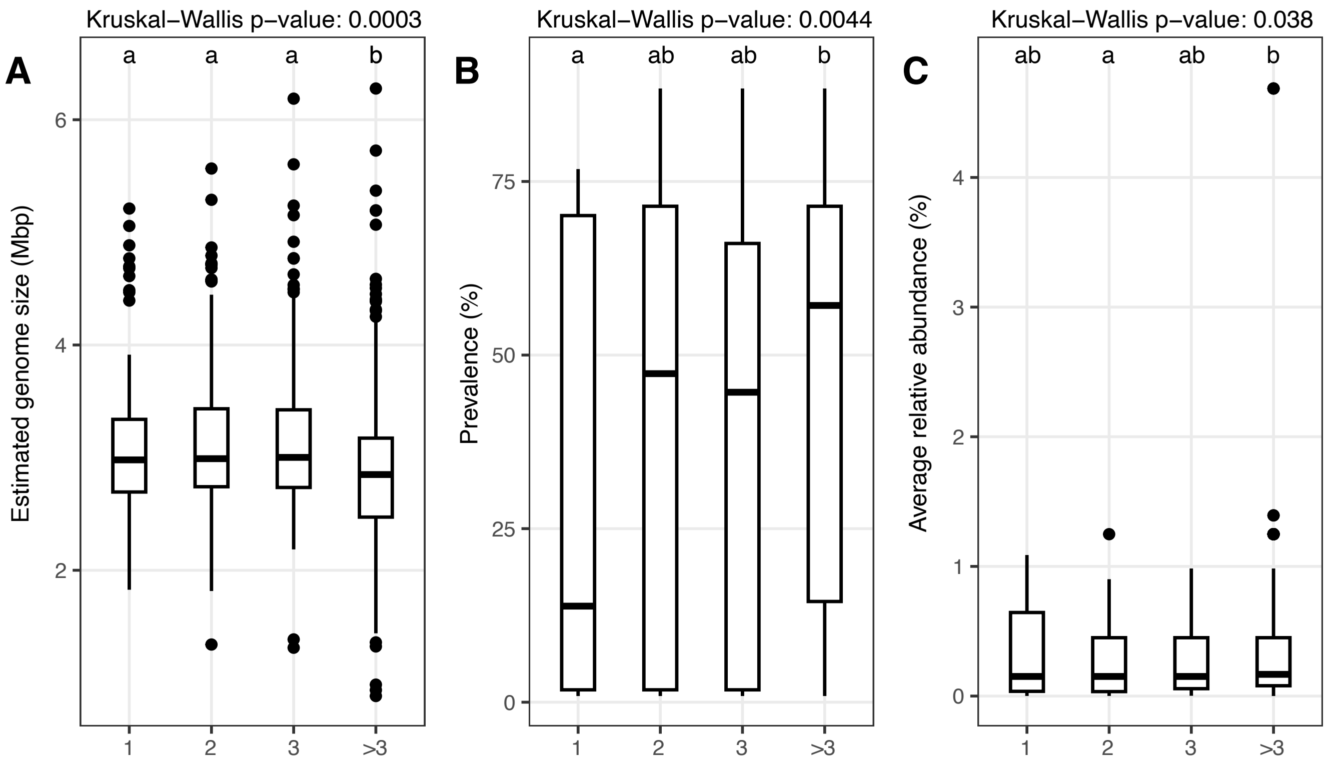


**Figure S7.** Boxplots comparing the (A) estimated genome size, (B) prevalence, and (C) average relative abundance of genomes from model communities (n = 527) that include 1, 2, 3, or more than 3 species per culture. Statistical significance was tested using the Kruskal-Wallis test ($p < 0.05$), followed by a Dunn's post-hoc test with Bonferroni correction. Groups with different letters are significantly different ($p < 0.05$).

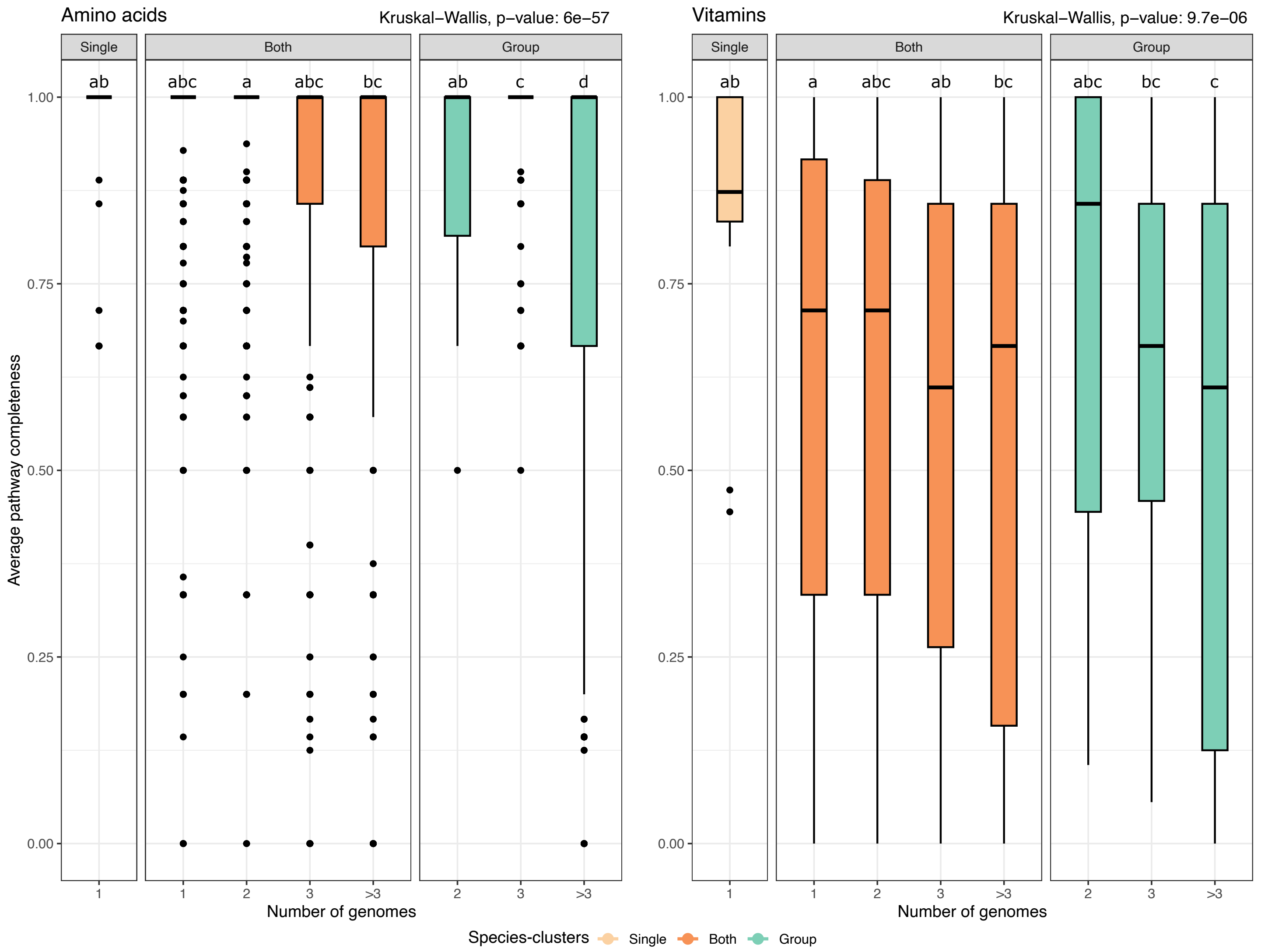


**Figure S8.** Pathway completeness of cultured genomes according to growth categories. Boxplots showing the average pathway completeness of all cultured high-quality genomes (> 90% completeness and < 5% contamination) for custom amino acid (A) and vitamin (B) modules, according to the number of genomes per culture, as well as the growth categories. Groups with different letters are significantly different (Kruskal-Wallis test followed by a Dunn's post-hoc test with Bonferroni correction, $p < 0.05$). Color coding displays whether the species cluster grew exclusively on their own (light orange), exclusively in groups (green), or both on their own as well as in groups (orange).

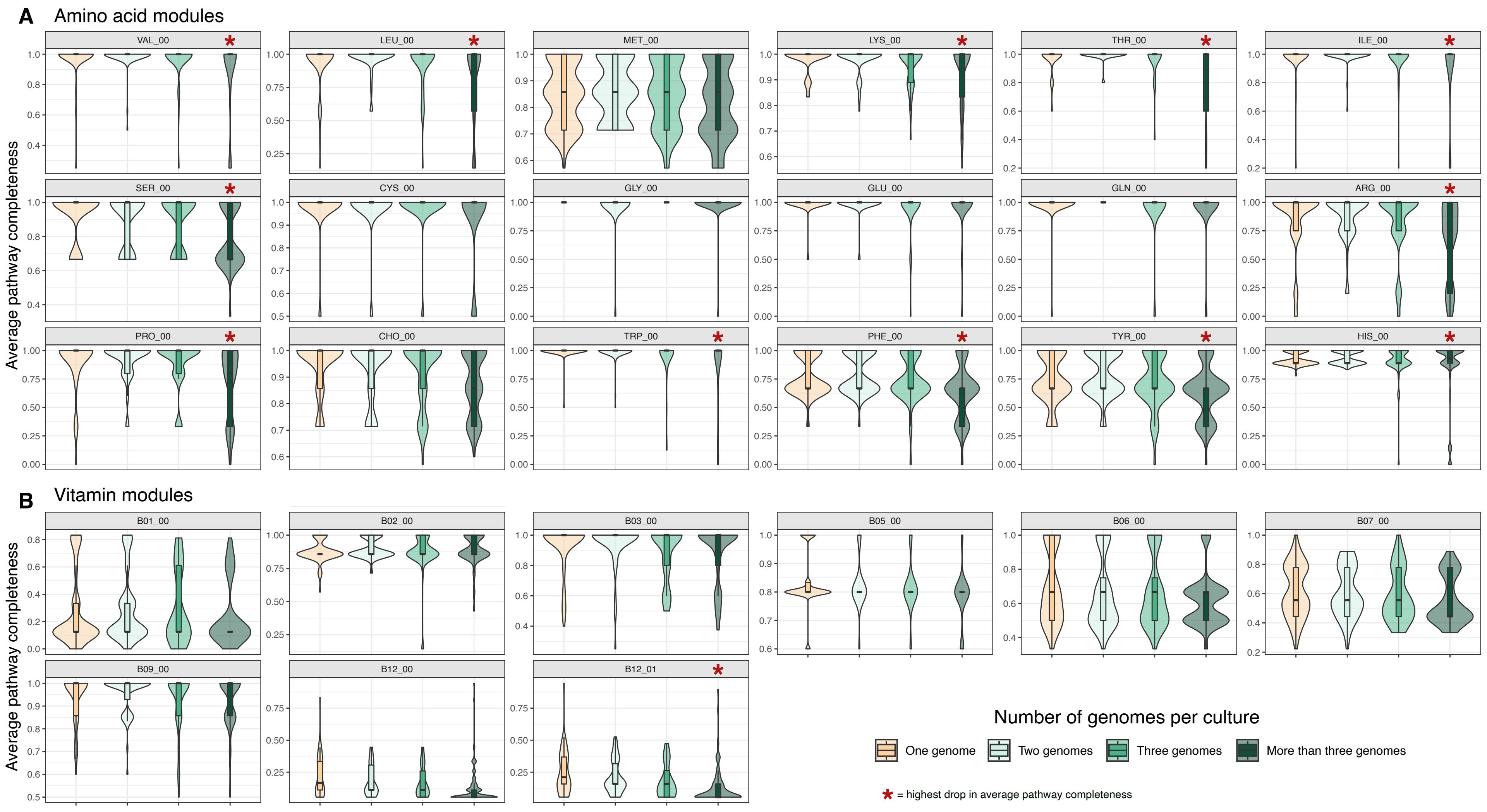


**Figure S9**. Violin plots showing the module completeness for the 305 high-quality genomes (>90% complete, <5% contaminated) from model communities and compared per community complexity: single-species (light orange), two-species (mint), three-species (green), and >3-species cultures (dark green). Red asterisks indicate modules with the largest relative decreases in completeness in >3-species communities compared to simpler communities.

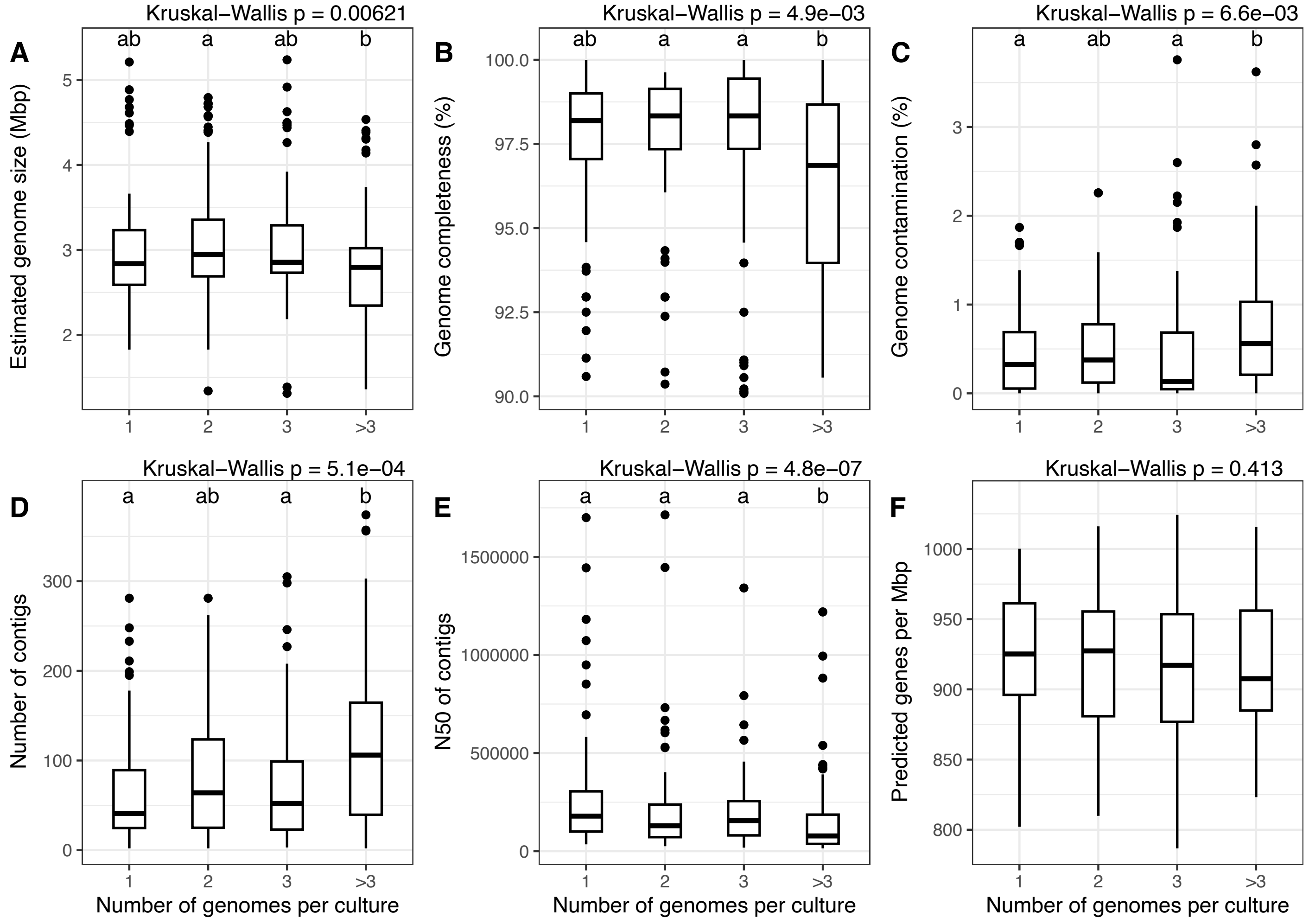


**Figure S10**. Box plots showing the comparison between high-quality cultivated genomes (n = 305) of various genomic metrics per community complexity (1, 2, 3, and more than three species per culture). These metrics include: (A) estimated genome size, (B) genome completeness, (C) genome contamination, (D) number of contigs per genome, (E) N50 of contigs, and (F) predicted genes per Mbp. Statistical significance was tested using the Kruskal-Wallis test ($p < 0.05$), followed by a Dunn's post-hoc test with Bonferroni correction. Groups with different letters are significantly different ($p < 0.05$).

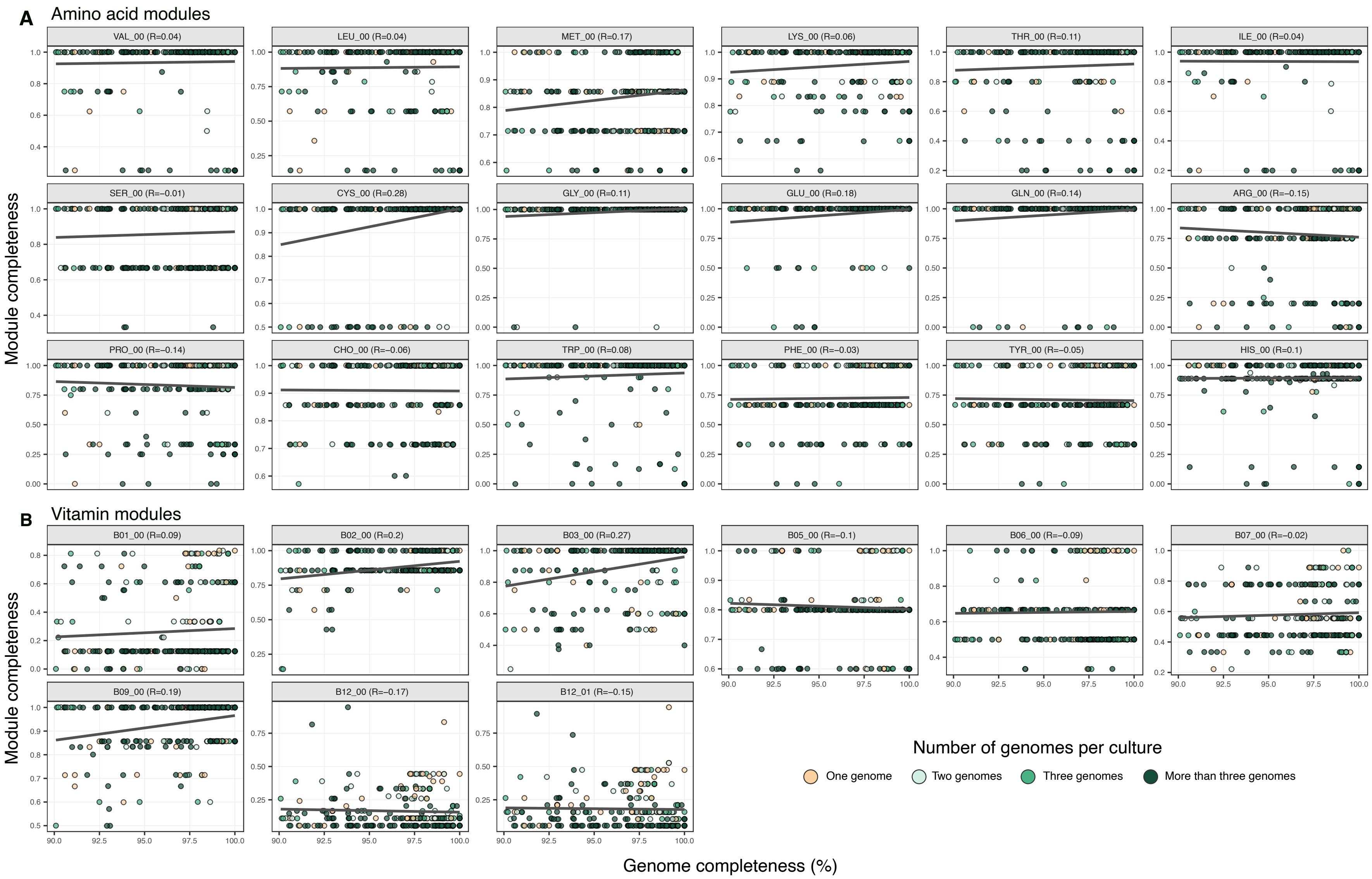


**Figure S11**. Scatter plots showing the relationship between genome completeness (x-axis) and module completeness (y-axis) for 305 high-quality genomes (>90% complete, <5% contaminated). Each point represents one genome and is color-coded by community complexity group: single-species (light orange), two-species (mint), three-species (green), and >3-species cultures (dark green). Linear trends (gray lines) were fitted, and Spearman's correlation values are indicated for each module. Most modules show weak correlations (R < 0.3), indicating that variations in biosynthetic potential are largely independent of genome completeness.

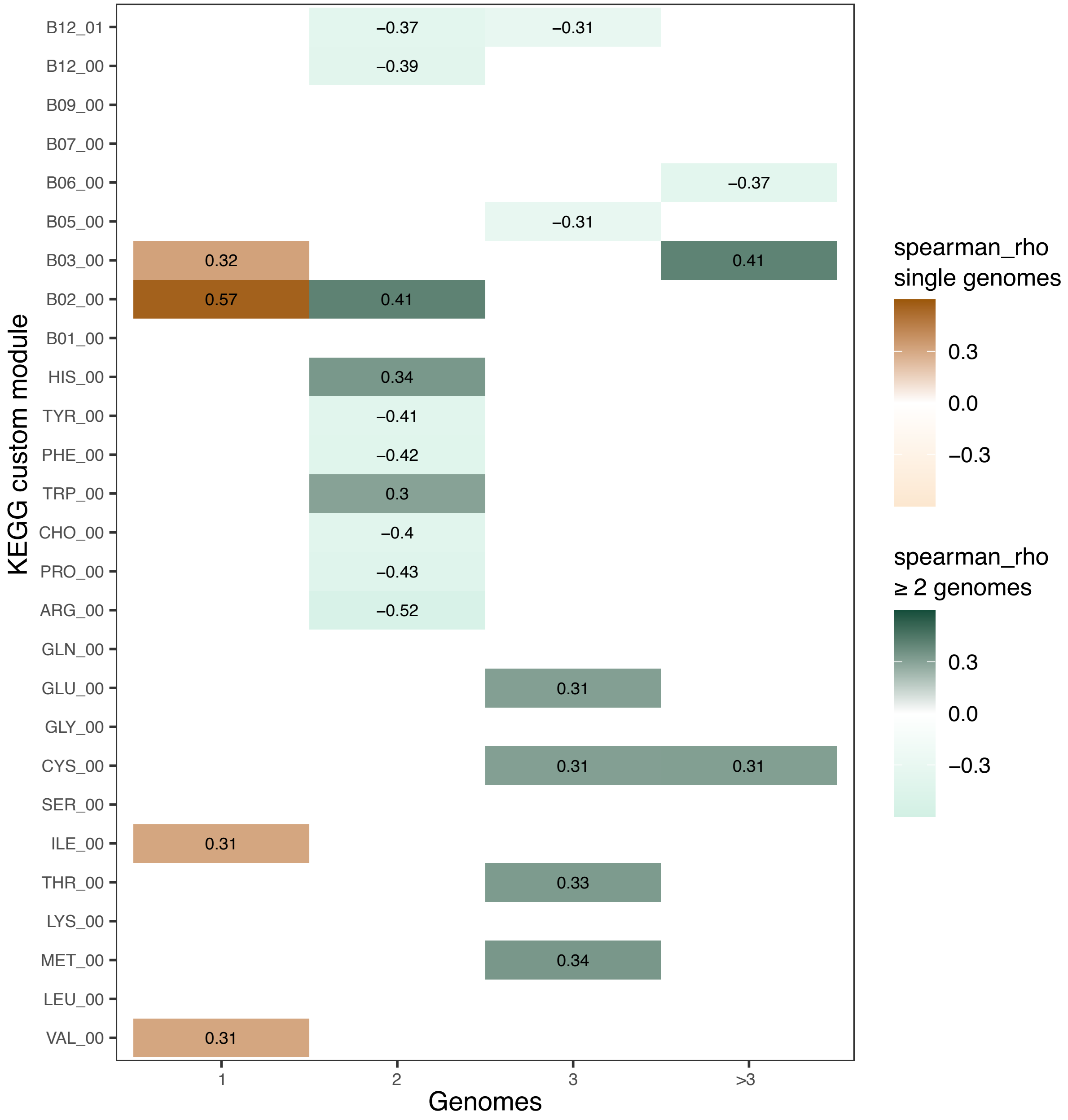


**Figure S12**. Heatmap showing Spearman's correlation values between genome completeness and module completeness for each biosynthetic module (rows) and community complexity group (columns). Only moderate correlations (R ≈ 0.3–0.5) are shown, with positive correlations in dark orange or green gradients for single genomes, and ≥ genomes per culture, respectively. Modules with no strong correlation are in white.

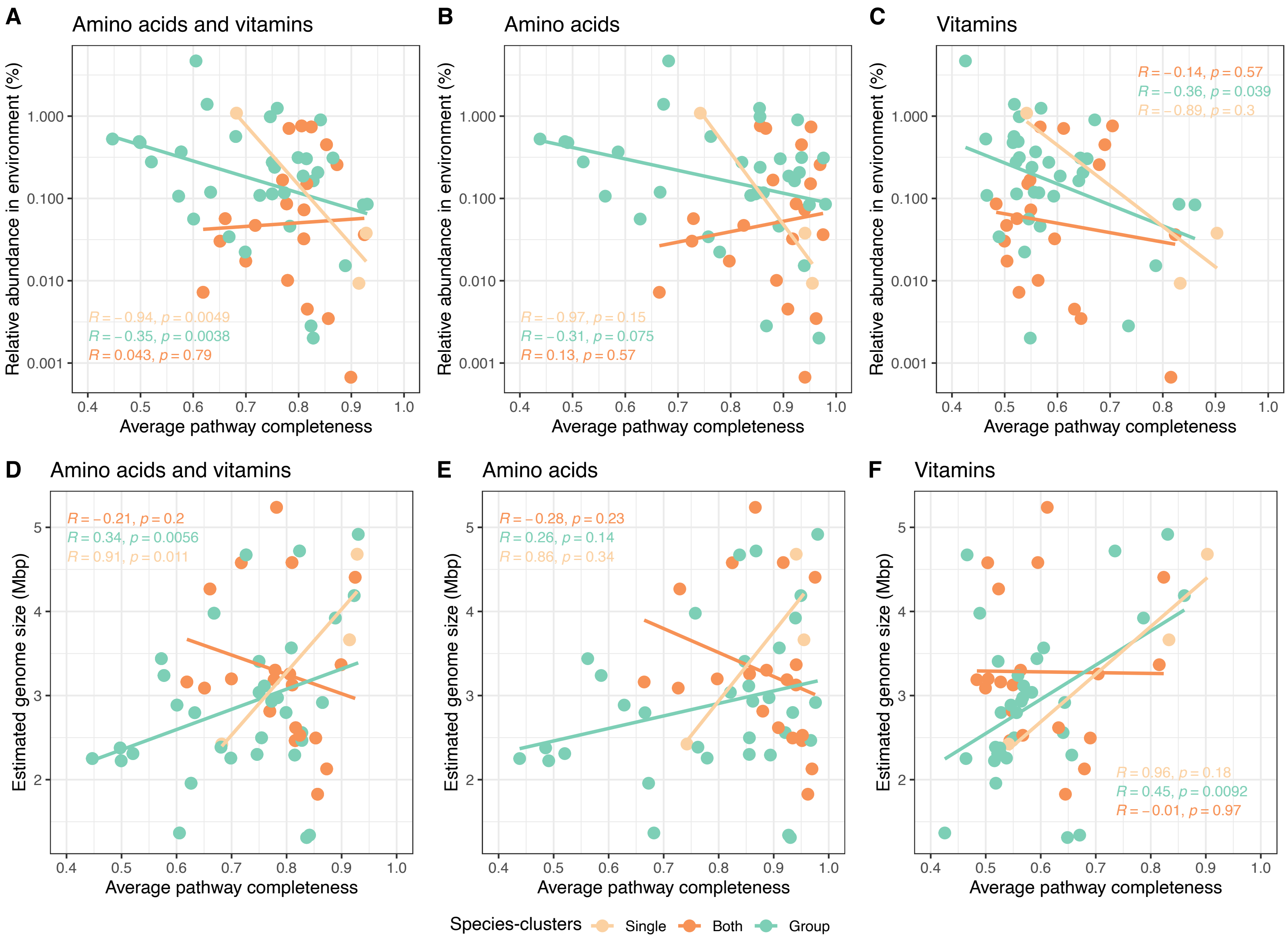


**Figure S13.** Relationship between pathway completeness, relative abundance, and genome size of cultured species-clusters. Overview of the correlation between average biosynthesis pathway completeness of cultured species-clusters and the relative abundance of these species-clusters in environmental samples as well as estimated genome size for three metabolic categories: Amino acids and vitamins (A, D), amino acids (B, E) and vitamins (C, F). Genomes with < 90% genome completeness and > 5% contamination were filtered out. Each data point represents one species-cluster, which either grew exclusively on their own (light orange), exclusively in groups (green), or both on their own as well as in groups (orange). Linear regression models were fitted to the data.

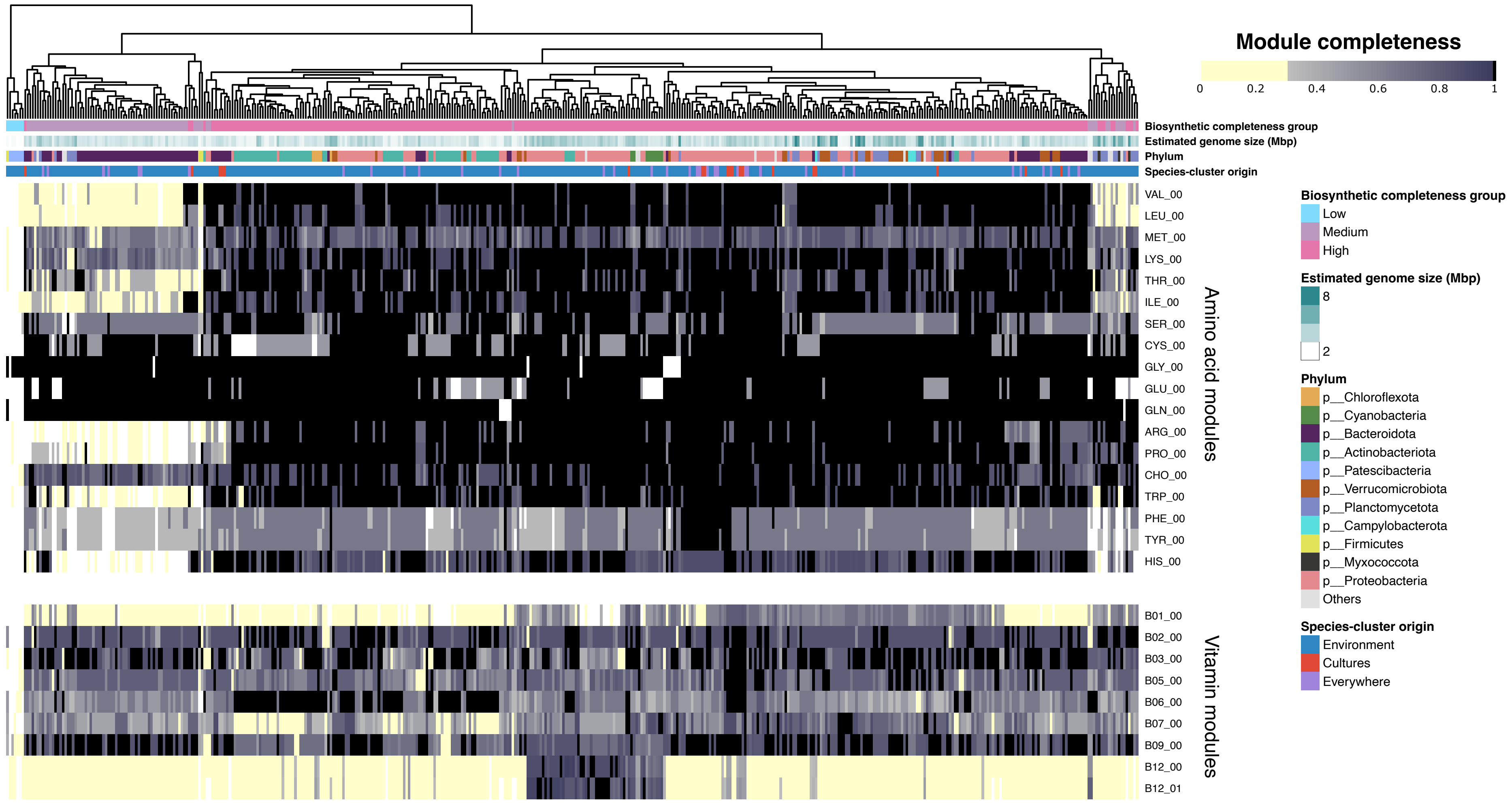


**Figure S14**. Figure S11. Biosynthetic potential clustering of the BalticMAG species catalog. Heatmap showing the pathway completeness of amino acid and vitamin biosynthesis pathways (based on KEGG custom modules) across high-quality species-cluster representative genomes (n = 450). The color gradient represents pathway completeness, with white = 0%, yellow = >0–30%, and black = 100% completeness. Genome annotations include: Biosynthetic completeness group (Low = light blue, Medium = light purple, High = pink), estimated genome size (white to dark teal gradient), taxonomic affiliation (color-coded by phylum), and genomic origin (environment-only = blue, culture-only = red, both = purple).

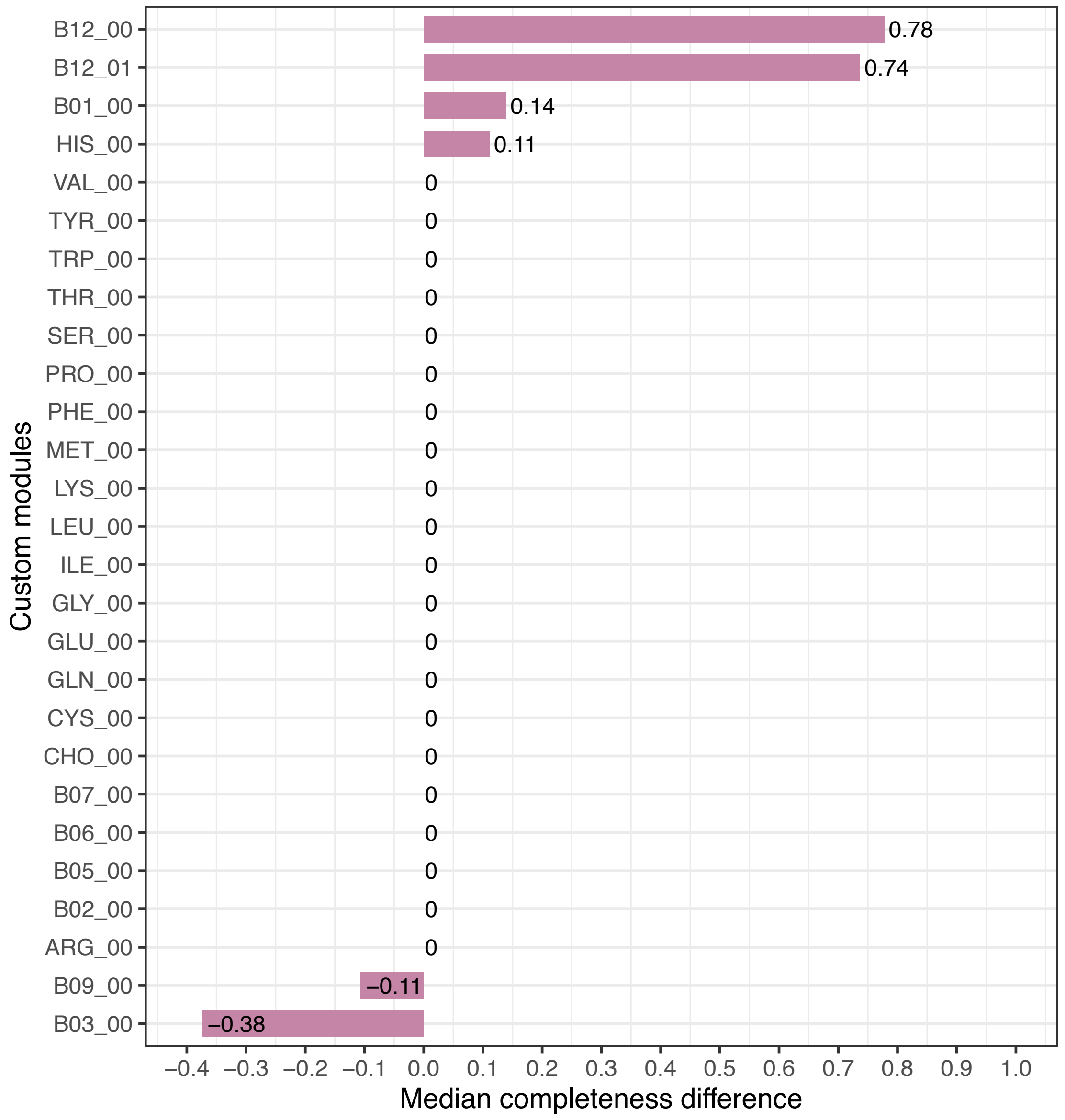


**Figure S15**. Median completeness differences for KEGG custom biosynthesis modules between Cluster A and Cluster B genomes from the High biosynthetic group. Positive values indicate modules are more complete in Cluster A, negative values indicate modules are more complete in Cluster B, and zero values show no differences in the median completeness between the two clusters.
